# The essential co-chaperone Sgt1 regulates client dwell time in the Hsp90 chaperone cycle

**DOI:** 10.1101/2025.04.24.648904

**Authors:** Sonja Engler, Florent Delhommel, Christopher Dodt, Abraham Lopez, Ofrah Faust, Valeria Napolitano, Grzegorz M. Popowicz, Rina Rosenzweig, Michael Sattler, Johannes Buchner

## Abstract

The Hsp90 machinery is the most complex chaperone system in the eukaryotic cell. It is characterized by numerous co-chaperones that modulate the function of Hsp90. In *S. cerevisiae*, most of these cofactors can be deleted without affecting viability. Of the three essential ones, only the function of Sgt1 remained enigmatic. Our *in vivo* and *in vitro* experiments define key structural elements and determine the essential function of Sgt1 in the chaperoning of client proteins. We show that yeast Sgt1 exhibits a unique binding mode to Hsp90. The simultaneous interaction of Sgt1 with Hsp90 and client proteins enhances client maturation efficiency. Specifically, Sgt1 stabilizes Hsp90-client complexes and prevents their dissociation by the co-chaperone Aha1. Together, our findings reveal a distinct regulatory mechanism of the Hsp90 function, highlighting Sgt1 as a critical modulator of chaperone cycle progression.

## INTRODUCTION

Cells rely on sophisticated folding machinery to deal with the constant risk of protein misfolding and aggregation.^1^ A key player in this machinery is the Heat Shock Protein 90 (Hsp90), one of the most abundant molecular chaperones in the cytosol of eukaryotic cells. It facilitates the proper folding and maturation of numerous ‘client’ proteins, including kinases, transcription factors, signaling proteins, and cell cycle regulators.^2,3^ By stabilizing these diverse clients, Hsp90 is crucial in supporting cellular signaling, growth, and stress responses.^4,5^

Hsp90 consists of three domains: an N-terminal domain (NTD) linked to a middle domain (MD) via a long linker and followed by a C-terminal domain (CTD). Dimerization *via* the CTD results in an open, V-shaped dimer.^6,7^ Driven by ATP-binding, Hsp90 undergoes large conformational changes from the open to closed states, including the dimerization of the two NTDs (closed I state), followed by the association of the NTDs with the MDs (closed II state).^7–9^ These transitions are essential for Hsp90’s function and are the rate-limiting step for ATP hydrolysis. The Hsp90 system represents the most complex chaperone network in the eukaryotic cytosol. The Hsp90 homolog of *S. cerevisiae* (Hsp82, referred to as Hsp90 for simplicity’s sake) works together with more than ten co-chaperones modulating the conformational processing of client proteins.^10,11^ With the exception of Hch1, all co-chaperones are conserved from yeast to humans. These co-chaperones contribute to different steps of the client folding process facilitated by Hsp90.

One of the most potent modulators of the conformational cycle of Hsp90 is the ATPase homolog 1 (Aha1). This non-essential co-chaperone accelerates the ATP hydrolysis of Hsp90 more than ten-fold.^12–16^ While early studies reported downregulation of client proteins upon Aha1 inhibition in yeast^14,17^, more recent reports consistently show increased activity and stability of diverse Hsp90 clients.^15,18–22^ Due to overlapping binding surfaces at the Hsp90-MD, Aha1 and some client proteins seem to compete for binding to Hsp90.^12,23,24^ These findings suggest that Aha1 may promote client release from Hsp90, effectively ‘clearing’ the chaperone for subsequent cycles.^25^

In yeast, only three co-chaperones are essential, one of them is the ‘Suppressor of the G2 Allele of Skp1’ (Sgt1). Sgt1 is conserved across all eukaryotes, with higher eukaryotes possessing two isoforms.^26–29^ Sgt1 has been proposed to be important for various cellular functions, including assembling the kinetochore and SCF E3 ubiquitin complex in yeast ^26,30,31^ and regulating innate immune systems in plants and animals.^32–35^ Importantly, Sgt1 appears to have a general impact on Hsp90-dependent client maturation as its knockdown in yeast affected all clients tested.^18^ In humans, Sgt1 is overexpressed in several cancers and in the brain of Parkinson’s disease patients.^36–39^ Structurally, Sgt1 comprises an N-terminal TPR (tetratricopeptide repeat) domain, a CS (chord-containing proteins and Sgt1) domain, and a C-terminal SGS (Sgt1-specific) domain. In yeast, Sgt1 forms dimers through the interaction of the TPR domain (Figure 1A). Both the TPR domain and the SGS domain have been reported to interact with structurally diverse client proteins.^30,40–43^ While the CS and SGS domains are conserved through evolution, the TPR domain is lacking in several species, including nematodes and insects.^44,45^ Unlike other Hsp90 co-chaperones, the TPR domain does not seem to mediate the interaction with the C-terminal tail of Hsp90. Instead, the CS domain of Sgt1 was reported to directly interact with the NTD of Hsp90 in plants.^40,46,47^ The unique SGS domain of Sgt1 is thought to be largely unstructured with small regions of helical propensity, but a defined structure is lacking.^47^ Notably, the human and plant SGS domain was reported to bind to the molecular chaperone Hsp70 rather than Hsp90, potentially linking the two chaperone systems.^48,49^

**Figure 1.**
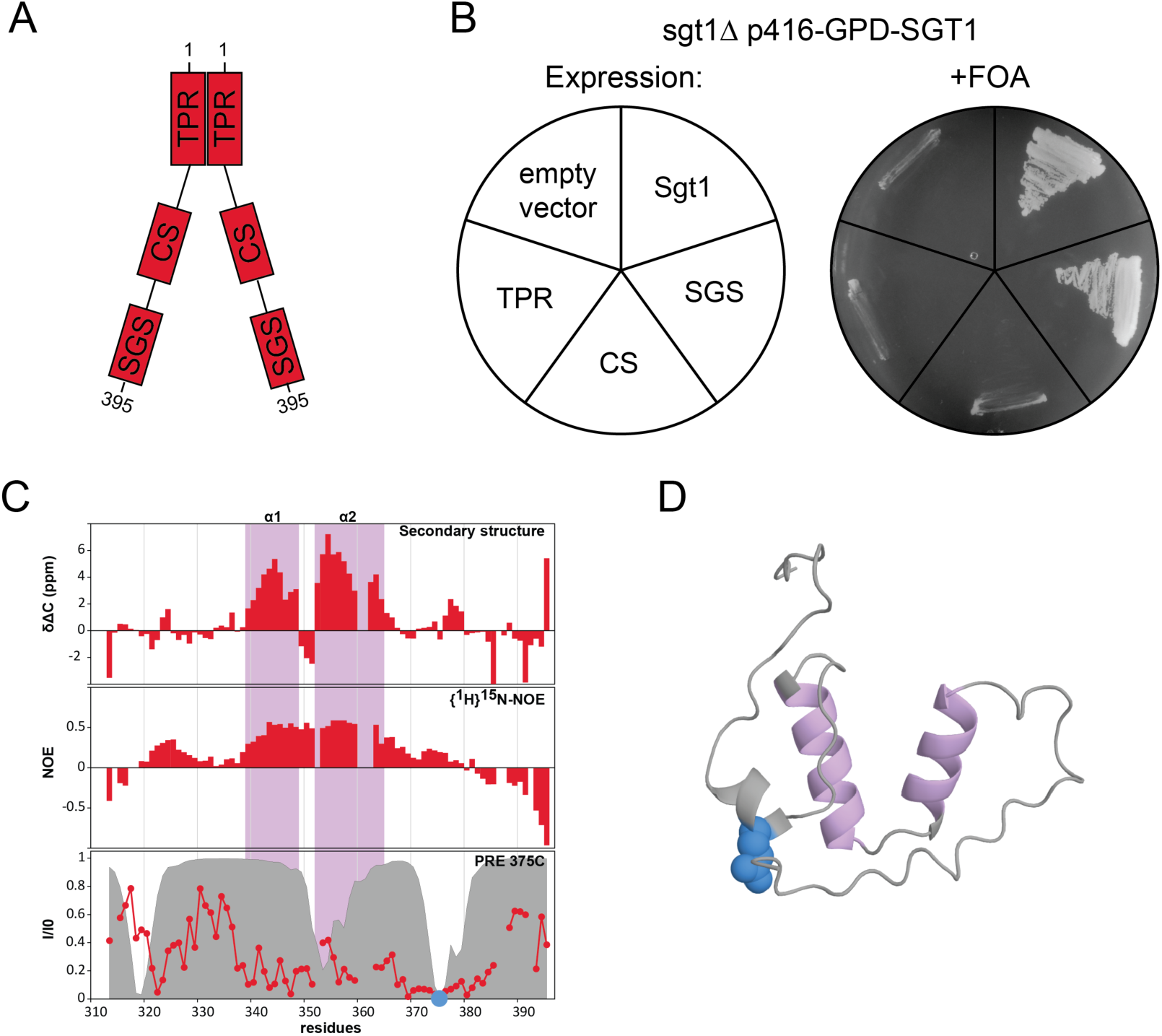
The SGS domain of Sgt1 performs the essential function. A) Schematic domain architecture of Sgt1. B) Determination of the essentail domain of Sgt1. Plasmid shuffling (right panel) of the BY4741 sgt1δ p416-GPD-SGT1 strain transformed with p415-GPD plasmids containing Sgt1, Sgt1 domains or empty vector as indicated in the left panel. C) The SGS domain of Sgt1 contains two stable helices, but without clear structures. Diagrams showing helical propensity (top), {1H}-15N NOE (middle), and PRE (bottom) measured with a probe introduced at position 375 (red line) compared to the predicted PRE, based on the AF2 model (gray volume). D) The AF2 model does not represent well the SGS structure. AF2 model showing in pink the regions of high helical propensity determined by NMR and the position of the PRE probe in blue.

Despite its importance, the specific function and underlying mechanism of Sgt1 within the Hsp90 cycle remain unclear. Here, we set out to define the structure-function relationship of Sgt1 and elucidate its function in the Hsp90 chaperone machinery through *in vitro* and *in vivo* experiments. Our results indicate that a specific structural element in the SGS domain is critical for its essential function, which is connected to the chaperone cycle of Hsp90. We show that Sgt1 facilitates client maturation by a unique mechanism. Together, our results show the complex interplay of the Hsp90 co-chaperone network to achieve client folding.

## RESULTS

### The SGS domain of Sgt1 is essential

Sgt1 is essential for yeast viability.^26^ To define the domain(s) of Sgt1 that harbor the essential function, we performed plasmid shuffling experiments, which allowed us to express Sgt1 domain constructs in a sgt1δ knockout strain and to determine their ability to support yeast growth. As expected, the expression of full-length Sgt1 from the plasmid rescued the chromosomal deletion of Sgt1. However, yeast cells expressing only the TPR, the CS, or both domains were inviable (Figure 1B). Remarkably, the expression of the SGS domain alone was sufficient to support wild-type-like growth, demonstrating that this domain harbors the essential *in vivo* function of Sgt1 (Figure 1B). Moreover, the SGS domain containing constructs, i.e., CS-SGS or TPR-L-SGS, in which a poly-GS linker connects the TPR and SGS domains, also showed wild-type-like growth (Figure S1A).

While structures of the CS and TPR domains of Sgt1 are known, the conformation of the unique SGS domain is unknown and has been predicted to be primarily disordered without a globular structure.^46,47^ We used NMR spectroscopy to characterize the conformation of the SGS domain. Secondary structure propensity derived from secondary ^13^C chemical shifts identified two helical regions between amino acids 339-348 (α1) and 352-364 (α2) (Figure 1C), in agreement with an AlphaFold 2 model. The AlphaFold model of the Sgt1-SGS predicts a defined conformation for the region comprising these two α-helices and a C-terminal loop. In this part of the model, the pLDDT scores are high, indicating a high level of confidence in the local geometry. Notably, the C-terminal loop appears to engage in stabilizing interactions with the second helix, suggesting a backbinding arrangement that could help to stabilize this segment. However, backbone flexibility assessed by heteronuclear {^1^H}-^15^N NOE analysis shows that the SGS domain is overall largely flexible but that the two helical regions are more rigid, with values around 0.5, still indicating significant conformational dynamics compared to what would be expected for a rigid globular domain. Consistent with this, paramagnetic relaxation enhancement (PRE) measurements on the SGS with a spin label probe attached to residue 375 (within the C-terminal loop) show significant line-broadening covering the helical regions but do not agree with theoretical PRE effects predicted for the AlphaFold model (Figure 1D). Similarly, experimentally determined residual dipolar couplings (RDC) do not support the AlphaFold model and are consistent with significant dynamic averaging of local structural elements (Figure S1B). These results suggest that the helical regions in the SGS domain alternate between conformational states of varying compaction without making stable long-range contacts. In summary, our results revealed that the SGS domain is the essential domain of Sgt1, and that this structure is a dynamic conformational ensemble with two helical regions.

### Sgt1 promotes GR maturation *in vivo* via its essential structural element

Next, we wondered whether the helical regions, which are evolutionary conserved (Figure S1C), contribute to the essential function of the SGS domain. To test this, we introduced proline residues in the first (F343P) or the second (M358P) helix to perturb the helical conformation and performed plasmid shuffling with these variants. While mutation of the first helix did not affect yeast cell growth, breakage of the second helix was lethal, demonstrating that the essential function of Sgt1 depends on the second helix (Figure 2A, B). To test whether the regions adjacent to the second helix are involved in the essential function, we mutated the linker region between the two helices to alanine (349-352Ala). Yeast cells expressing only the mutated Sgt1 showed no growth defect, indicating that the linker between the two helices is not essential (Figure S2A). Truncation of the C-terminal tail revealed that a deletion up to amino acid 372 induced reduced but robust growth. In contrast, a truncation up to amino acid 371 was lethal (Figure 2A. Figure S2B). Interestingly, the second helix ends at amino acid 364. Thus, this experiment identified the second helix plus the eight subsequent amino acids of the C-terminal tail as critical for the essential function of Sgt1 (Figure 2B). The comparison of NMR spectra from the corresponding truncated constructs revealed a severe line broadening in CS-371 compared to CS-372, with only negligible changes in chemical shifts. This likely results from a pronounced reduction in the observable population of the helical conformation and an increase in intermediate exchange (Figure S2C, D), indicating the stability of the helices in the SGS as critical for Sgt1’s function.

**Figure 2.**
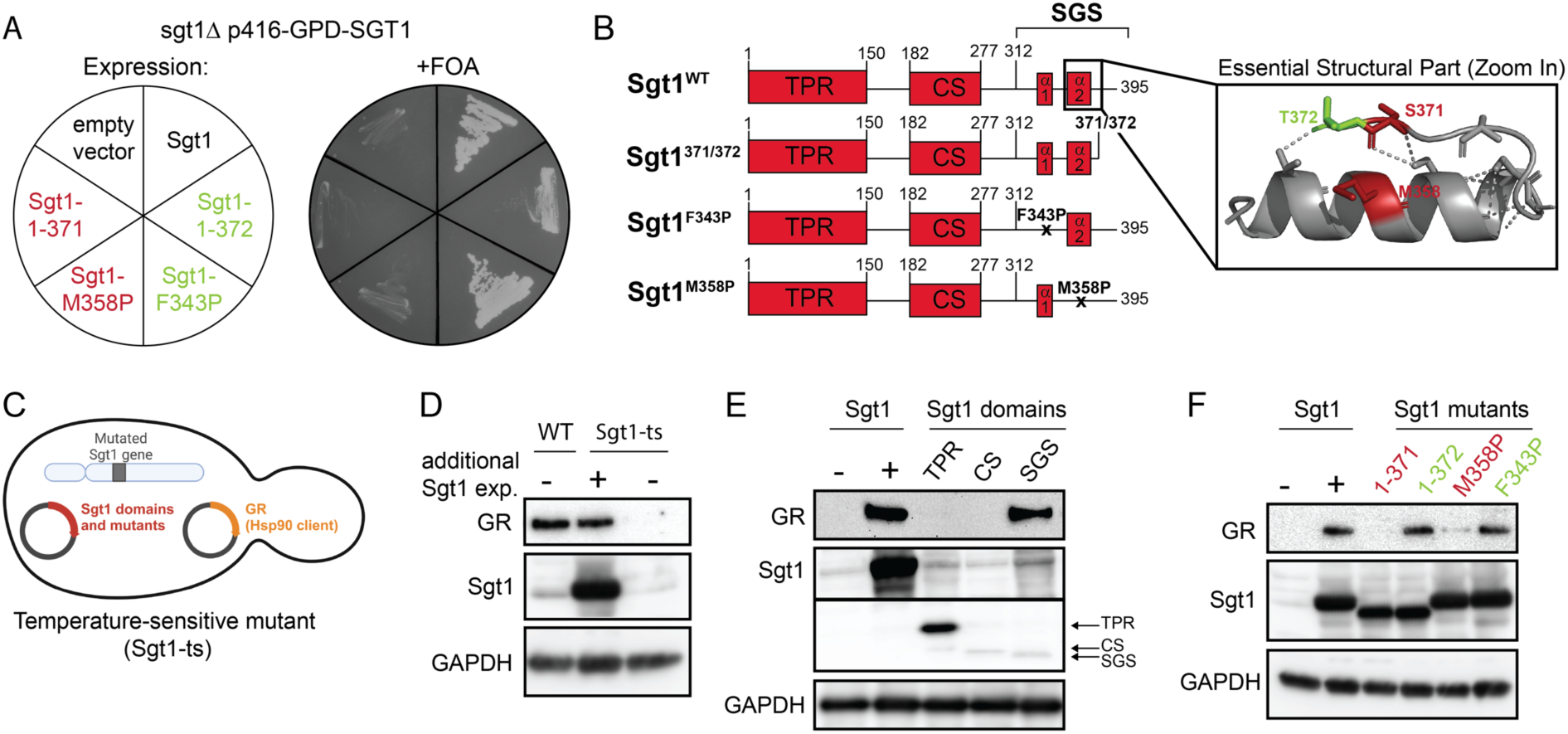
Sgt1 stabilizes GR *in vivo*. A) In vivo analysis of the essential structural element of Sgt1. Plasmid shuffling (right panel) of the BY4741 sgt1δ p416-GPD-SGT1 strain transformed with p415-GPD plasmids containing Sgt1, Sgt1 mutants or empty vector as indicated in the left panel. B) Schematic domain architecture of Sgt1 truncations and point mutants, with a zoom in of the essential structural element of the AF2 structure. The key residues involved in the viability of the mutants are highlighted. C) Schematic illustration of the temperature-sensitive strain Sgt1-3 (Sgt1-ts). The strain harbors three mutations in the SGT1 gene, resulting in a destabilized, non-functional Sgt1 protein at non-permissive temperatures (30°C). The strain was transformed with a plasmid for Sgt1 expression (including single domains and mutants) along with a plasmid for expression of GR. D) Effect of Sgt1 on GR levels. The yeast strain BY4741 (WT) and Sgt1-ts expressing Sgt1 (+) or empty vector (-) were grown at 30°C. The levels of GR and Sgt1 were examined by immunoblot. GAPDH was used as a loading control. A representative immunoblot from three individual experiments is shown. E) Analysis of the essential structural element of Sgt1 for chaperoning. The yeast strain Sgt1-ts expressing Sgt1 (+), empty vector (-) or Sgt1 domains (TPR, CS and SGS) was grown at 30°C. GR and Sgt1 levels were examined by immunoblot. GAPDH was used as a loading control. A representative immunoblot from three individual experiments is shown. F) The yeast strain Sgt1-ts expressing Sgt1 (+), empty vector (-) or Sgt1 mutants (1-371, 1-372, M358P and F343P) was grown at 30°C. GR and Sgt1 levels were examined by immunoblot. GAPDH was used as a loading control. Lethality of the mutants 1-371 and M358P is indicated in red, viability of the mutants 1-372 and F343P is indicated in green. A representative immunoblot from three individual experiments is shown.

To study the chaperone function of Sgt1 *in vivo*, we used the temperature-sensitive strain Sgt1-3 (hereafter referred to as Sgt1-ts). Since a complete knockout of Sgt1 is not viable, the ts-strain provides an alternative for studying both lethal and viable Sgt1 mutations. This strain harbors multiple mutations in the SGT1 gene, resulting in a dysfunctional Sgt1 protein at non-permissive temperatures.^26^ We transformed the strain with plasmids, allowing the simultaneous expression of Sgt1 (full-length, domains, and mutants) along with the glucocorticoid receptor (GR) (Figure 2C). GR is a well-studied transcription factor that is strongly dependent on the Hsp70 and Hsp90 chaperone system for its maturation and activation.^18,50–52^ At non-permissive temperatures, GR levels were severely reduced in the Sgt1-ts strain compared to the WT (Figure 2D). Reconstitution of Sgt1 expression in the ts-strain restored GR levels to those observed in WT yeast, indicating that the stability of GR is strongly dependent on Sgt1 in yeast. To test how the influence of Sgt1 on GR maturation relates to its essential function in yeast, we determined to what extent the different structural elements of Sgt1 are involved in this process *in vivo*. To this end, we expressed the individual Sgt1 domains in the ts-strain and analyzed GR levels. We could not detect GR when we expressed only the TPR or CS domain. However, in the presence of the SGS domain, GR accumulated to WT levels (Figure 2E). We used the ts-strain to further define the requirements for GR chaperoning compared to the essential function of Sgt1. We expressed Sgt1 constructs that either did not support yeast growth in the Sgt1 deletion strain (1-371 and M358P) or which were able to perform the essential function of Sgt1 (1-372 and F343P). The analysis of GR levels showed that the viable mutants stabilized GR while the lethal mutations did not (Figure 2F). Thus, the minimal structural element that carries out the essential function of Sgt1 is also responsible for chaperoning GR. In conclusion, these results suggest that Sgt1’s function in the context of chaperoning proteins is essential *in vivo*.

### Sgt1 binds Hsp90 via its CS and SGS domains

Many co-chaperones bind to Hsp90 through their TPR domains, and thus, it had been expected that Sgt1 would interact with Hsp90 similarly.^30^ However, structural studies revealed that in plants, the TPR domain of Sgt1 is not involved in Hsp90 binding.^46^ Analytical ultracentrifugation (aUC) analysis confirmed that to be true also for yeast Sgt1 and Hsp90. While aUC titration experiments using labeled Sgt1 and Hsp90 revealed a K_D_ of 0.51 µM, no interaction was detected between Sgt1-TPR domains and Hsp90 (Figure S3A,B). To gain further insight into the interaction of Sgt1 with Hsp90, we analyzed the interaction by NMR spectroscopy. Titrations of Sgt1-CS-SGS with isolated domains of Hsp90 confirmed the known binding site to the NTD^46^, with limited intensity reduction and chemical shift perturbations (CSP) seen for residues 180-210 (Figure 3A, S3E). Surprisingly, the addition of unlabeled Hsp90-MD induced a drastic intensity reduction of NMR signals in Sgt1-CS. Further NMR titrations revealed that the interactions of Sgt1 with the Hsp90-MD are stronger than with the NTD (Figure S3F). Consistent with this, aUC analysis using labeled Sgt1 and isolated Hsp90 domains confirmed strong binding to the MD, while no interaction with the NTD was detected, indicating a transient and possibly weak interaction (Figure S3C,D). To gain a better understanding of the Sgt1 binding site in the Hsp90-MD, we determined the crystal structure of the Sgt1-CS/Hsp90-MD complex (Figure 3B). The structure confirms the interfaces observed by NMR and shows that the CS domain binds from the edge of its β-sandwich fold to the top of the Hsp90-MD helix α2. Interestingly, we have previously identified a similar binding site on the Hsp90-MD for the Hsp90 co-chaperone NudC.^53^

**Figure 3.**
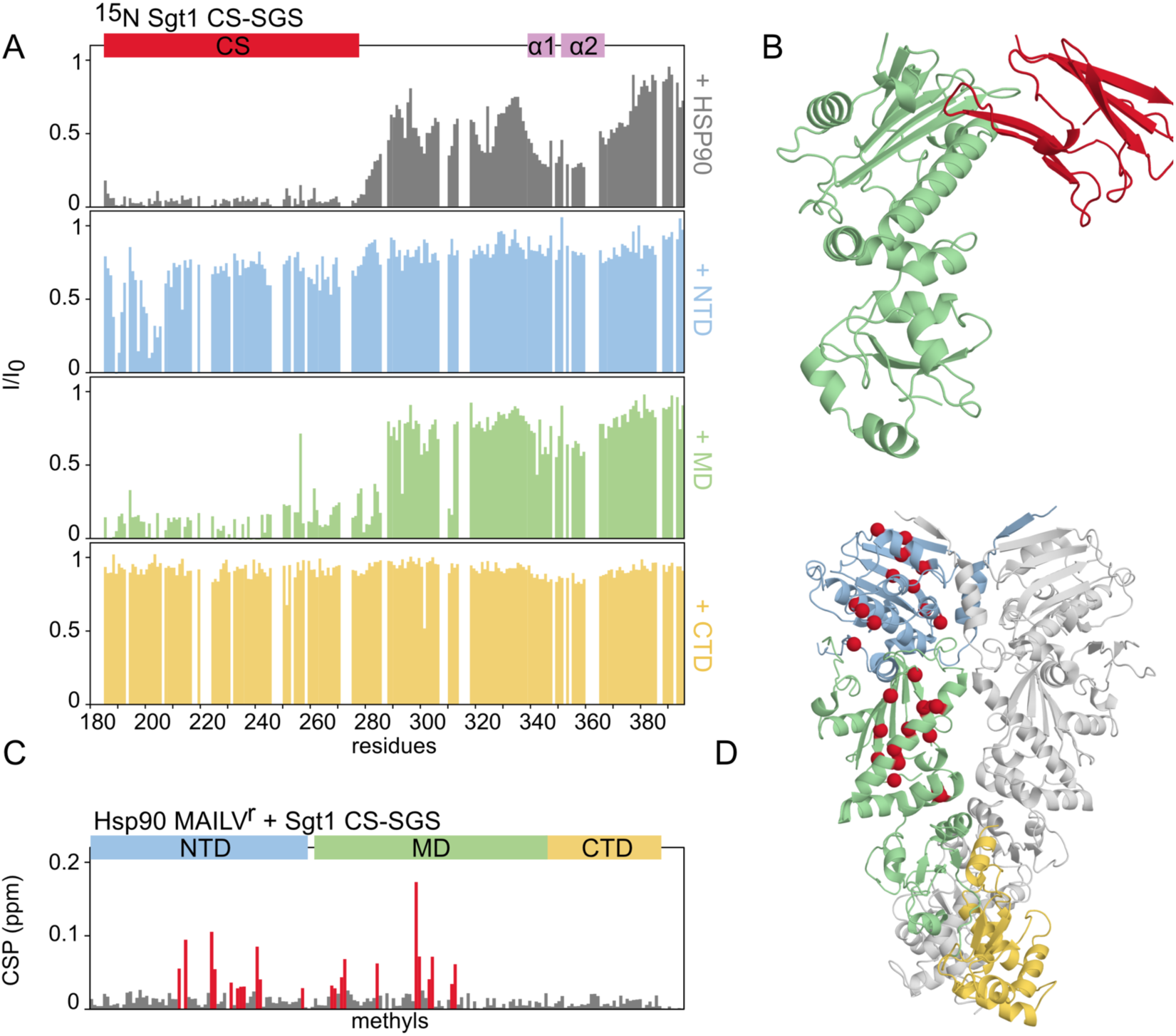
Sgt1-CS interacts with the NTD and MD of Hsp90. A) NMR analysis of the interaction of Sgt1 with Hsp90 domains. The relative intensity changes of the ^1^H-^15^N HSQC of the Sgt1-CS-SGS before and after the addition of an equimolar amount of Hsp90-FL (grey), NTD (blue), MD (green) and CTD (yellow) show the interactions of the Sgt1-CS domain with both NTD and MD. B) Crystal structure of the Hsp90-MD (green), Sgt1-CS complex (red). C) CSP plot of ^1^H-^13^C TROSY of the titration on Hsp90 methyl-labeled, with Sgt1-CS-SGS, also shows both interactions in the full-length context. D) Structure of Hsp90 reporting methyl CSP from plot C on the structure. Residues for which CSP>0.015 are shown in red. The perturbations fit well with the known interfaces of Sgt1-CS.

To learn more about this interaction, we performed the reversed titration, using full-length MAILV^r^ methyl-labeled Hsp90 and adding unlabeled Sgt1-CS-SGS. This titration validated the interaction observed with the isolated domains and showed that both the NTD and MD binding sites are occupied in the full-length context (Figure 3C, D). Titration of Sgt1-CS-SGS constructs to Hsp90-MD-CTD MAILV^r^ shows the expected MD binding site with small additional chemical shift changes in the CTD compared to those seen with the CS-only construct, suggesting that these were induced by the SGS region. These changes were not observed in the isolated domains and are overall consistent with a weak and dynamic interaction between Sgt1-SGS and Hsp90-CTD (S3G-I). Similar transient interactions with the client-binding region of the CTD involving flexible helical regions have been previously observed with other CS-containing co-chaperones, p23 and NudC.^53,54^

We were intrigued by the apparent paradox that the CS domain is the main contributor to the interaction with Hsp90, while the SGS domain is essential for Sgt1’s function. Since all constructs in the viability assays were expressed under the robust GPD promoter, we explored whether the SGS domain alone could support yeast growth using the native promotor (1000bp upstream of the SGT1 gene). Interestingly, under these conditions, the SGS domain was insufficient to maintain yeast viability. However, the expression of the CS-SGS construct did sustain yeast growth, indicating that the CS domain contributes an additional, albeit non-essential, function (Figure S3J). These findings suggest that the CS domain may provide a stable interaction with Hsp90, enabling the SGS domain to effectively perform its chaperone function. Altogether, our data identify a previously unknown binding site of Sgt1-CS in the Hsp90 MD. While the SGS domain exhibits a dynamic and weak interaction with the Hsp90 CTD, the interaction of Sgt1 and Hsp90 is primarily mediated by the CS domain. This raises the question of why the SGS domain is so critical for yeast viability.

### Sgt1 does not interact with the Hsp70 system

Previous work showed an interaction of the SGS domain of Sgt1 with Hsp70 and thus suggested that Sgt1 may function as an adaptor protein linking the Hsp70 and Hsp90 chaperone systems.^48,49^ aUC analysis with labeled Sgt1, however, did not reveal complex formation with Hsp70. NMR binding experiments with ^15^N-labeled CS-SGS further confirmed that Hsp70 does not interact with Sgt1 (Figure S4A, B). To test whether Sgt1-SGS might interact with JDP (referred to here as Hsp40) co-chaperones instead, as has been previously seen for the Hsp90 cofactor NudC, we tested the yeast Hsp40 protein Ydj1. The addition of a two-fold excess of Ydj1 to ^15^N-labeled CS-SGS showed only a very weak binding (Figure S4A), suggesting that Sgt1 does not form a substantial interaction with Ydj1. These results strongly indicate that Sgt1 does not directly link the Hsp70 and Hsp90 chaperone systems.

### Ternary complex formation by Sgt1, GR-LBD, and Hsp90 promotes client maturation efficiency

We next asked whether the SGS domain is required for interaction with client proteins, while the CS domain mediates Hsp90 binding. To test whether Sgt1 can directly interact with GR, we performed NMR analysis using the ^15^N-labeled CS-SGS fragment together with the ligand binding domain of GR (GR-LBD), which is targeted by chaperones within the GR.^55–59^ The NMR results showed line-broadening and chemical shift changes for SGS residues in the presence of the GR-LBD. Analysis of these changes suggests that the GR-LBD binds specifically to the helices within the SGS domain of Sgt1 (Figure 4A, S4C). aUC analysis using labeled GR-LBD additionally confirmed the complex formation of GR-LBD with Sgt1 (Figure 4B). As our results so far showed that Sgt1 can bind to both the GR-LBD and Hsp90, we wondered whether forming a ternary complex was possible. To address this question, we monitored the complex formation of the labeled GR-LBD with Hsp90 in the presence of Sgt1 by aUC and observed a peak at 9.3 S, indicative of the formation of a ternary complex of Sgt1 with the GR-LBD and Hsp90 (Figure 4B, S4D). Remarkably, the additional presence of Ydj1 and Hsp70 did not influence this complex formation, indicating a strong interaction and further supporting the conclusion that Sgt1 does not interact significantly with either component of the Hsp70 system (Figure S4E).

**Figure 4.**
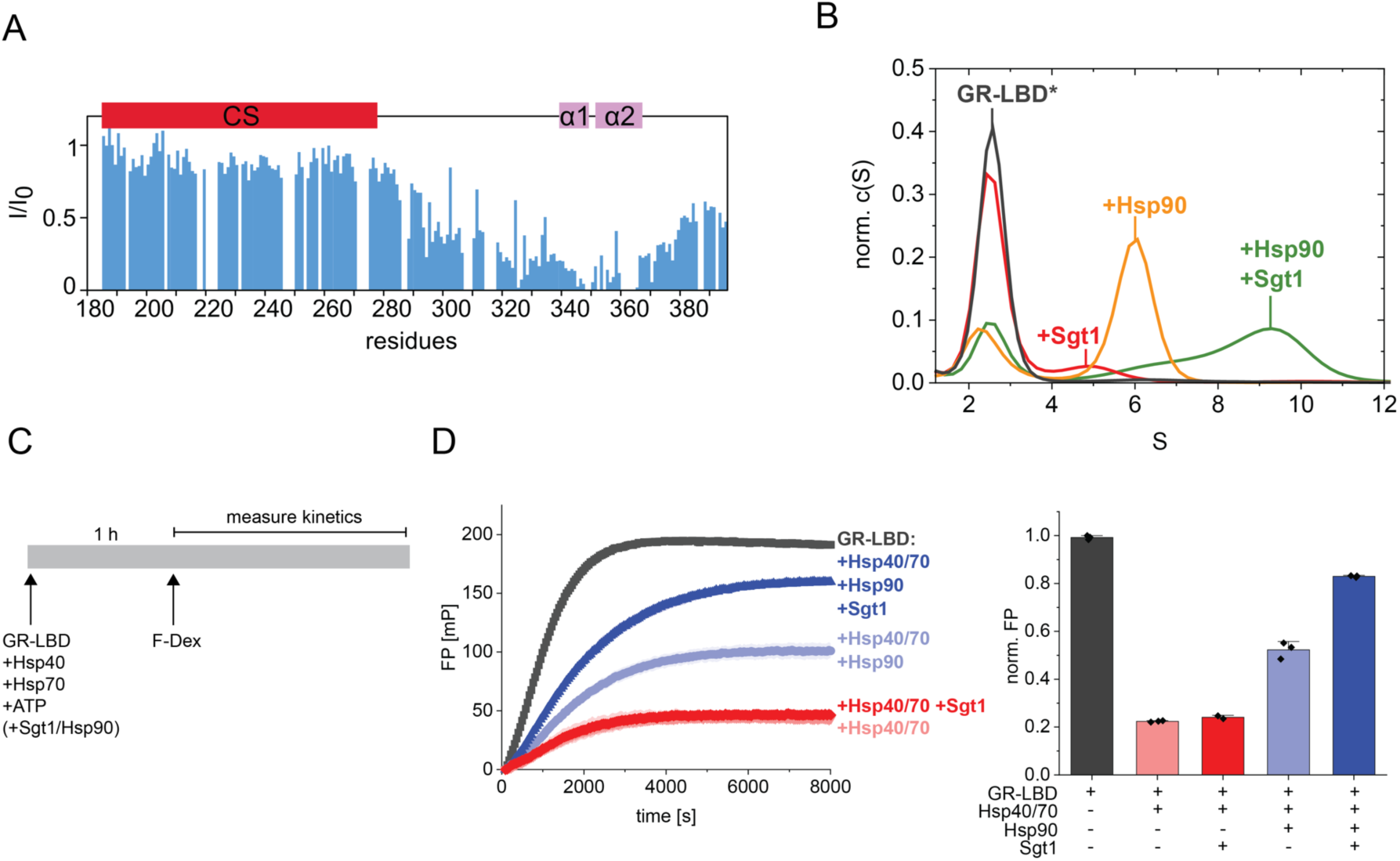
Sgt1 facilitates client folding *in vitro*. A) Relative intensity changes from the ^1^H-^15^N HSQC of Sgt1-CS-SGS before and after the addition of GR-LBDm shows that GR interacts with the SGS domain. B) Sgt1 forms a ternary complex with Hsp90 and GR-LBD. Complex formation analysis of labeled GR-LBD* with Sgt1 and Hsp90 by aUC sedimentation is shown. Normalized c(S) distributions were plotted against the sedimentation coefficient S. [GR-LBD*: 500 nM, Hsp90: 10 µM, Sgt1: 10 µM, ATP: 2mM] C) Experimental setup to follow hormone binding to apo-GR-LBD. GR-LBD (1 µM) was preincubated with Hsp40 (Ydj1, 2 µM), Hsp70 (12 µM), ATP (5 mM) and chaperones (Hsp90: 12 µM, Sgt1: 24 µM) for 1 h. The reaction was initiated by addition of fluorescein-dexamethasone (F-Dex, 100 nM). D) Effect of Sgt1 on Hsp90-induced hormone binding to GR-LBD. Fluorescence polarization measurement as described in C). Left panel: The binding kinetics of F-Dex to GR-LBD over time is shown, with the initial millipolarization (mP) value set to zero immediately after F-Dex addition. Right panel: Normalized change of fluorescence polarization over time. Results are presented as mean ± standard deviation.

Next, we sought to assess the influence of Sgt1 on client maturation by the Hsp90 chaperone system. To do this, we employed a fluorescence anisotropy-based assay that monitors the hormone-binding capacity of the GR-LBD. ^52,60,61^ This assay takes advantage of the fact that hormone binding to the GR-LBD is strongly dependent on its maturation state in the folding process. During the early stages of folding, GR-LBD is bound to the Hsp40/Hsp70 system, which keeps the client in an unfolded conformation, thereby preventing hormone binding. Upon release from Hsp70 and subsequent transfer to Hsp90, the GR-LBD can proceed through the folding pathway, ultimately regaining its hormone-binding ability.^60^ We incubated the GR-LBD with different combinations of the proteins Hsp40/Hsp70, Hsp90, and Sgt1 and monitored the binding of the fluorescence-labeled hormone dexamethasone (Figure 4C). The addition of Hsp40/Hsp70 to the GR-LBD significantly reduced its hormone-binding capacity. However, adding Sgt1 alone did not improve the hormone binding capacity (Figure 4D). As expected, adding Hsp90 increased the hormone binding capacity by around 30%. Remarkably, the simultaneous addition of Hsp90 and Sgt1 increased the hormone binding of GR-LBD by about 60%, indicating a synergistic effect of Hsp90 and Sgt1 in client maturation (Figure 4D). Together, our results suggest that Sgt1 is not sufficient to recruit the GR-LBD from the Hsp40/Hsp70 deadlock but that the direct interaction of Sgt1 with the GR-LBD and Hsp90 facilitates the formation of a client-Sgt1-Hsp90 complex in which Sgt1 enhances client folding by Hsp90.

### Sgt1 stabilizes the Hsp90-GR interaction and prevents client release by Aha1

Next, we investigated the interplay of Sgt1 with members of the Hsp90 co-chaperone system, focusing specifically on co-chaperones involved in client transfer and processing. We tested whether these co-chaperones can form complexes with Hsp90 and Sgt1. In line with previous results^40^, our aUC analysis revealed that Sti1 (hereafter referred to by the more commonly used name Hop), a key co-chaperone for client transfer, binds to both the Hsp90-Sgt1 and the GR-Hsp90-Sgt1 complexes (Figure S5A, B), indicating that Sgt1 adds an additional layer of regulation to the co-chaperone system. Sba1 (hereafter referred to as p23), a co-chaperone that selectively binds Hsp90 in its ATP-bound state and supports client maturation, was found to compete with Sgt1 for Hsp90 binding when present in great excess (Figure S5C,D). Consistent with earlier studies^40^, we found that Sgt1 does not affect Hsp90’s ATPase activity, even in the presence of p23 or Hop (Figure S5E).

Notably, for Aha1, a co-chaperone that accelerates the conformational cycle and binds to the Hsp90-MD ^14,24^, our data indicate a distinct interaction with Sgt1. aUC experiments revealed that labeled Sgt1 and Aha1 compete for binding to Hsp90 when no client protein is present (Figure 5A). The same result was obtained using labeled Aha1 (Figure 5B). In line with the comparable binding affinities of the Hsp90-Aha1 (K_D_ = 0.16 to 3.8 µM^12–15,62^) and the Hsp90-Sgt1 interaction (K_D_ = 0.51 µM), the two co-chaperones seem to be equally capable of releasing each other (Figure 5C).

**Figure 5.**
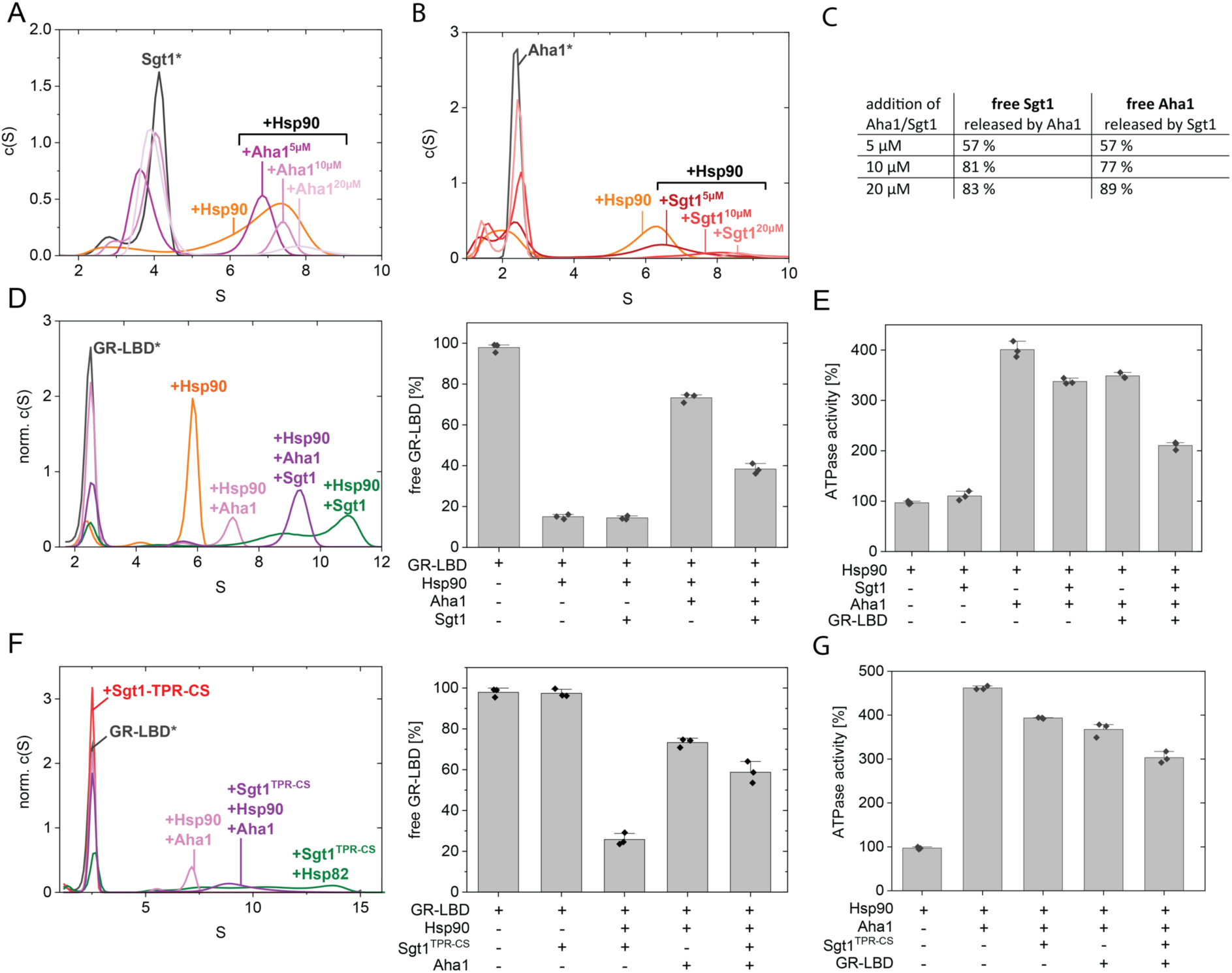
Sgt1 stabilizes the Hsp90-GR interaction and prevents Aha1-induced GR release from Hsp90. A) Competition between Aha1 and Sgt1 for Hsp90 binding. Release of Sgt1 from Hsp90 by Aha1. An aUC sedimentation analysis was performed using labeled Sgt1, Hsp90 and varying concentrations of Aha1. Normalized c(S) distributions were plotted against the sedimentation coefficient S. [Sgt1*: 500 nM, Hsp90: 5 µM, Aha1: as indicated, ATP: 5 mM] B) Release of Aha1 from Hsp90 by Sgt1. aUC sedimentation experiment as described in A), but using labeled Aha1 and varying concentrations of Sgt1. C) Quantification of free Sgt1 and free Aha1 from panels A) and B). The percentage of free Sgt1 (4 S) and free Aha1 (2.5 S) was determined by integrating the area under their respective peaks in the normalized c(S) distributions. D) Effect of Sgt1 on the Aha1-induced release of the GR-LBD from Hsp90. Left panel: Representative complex formation analysis of labeled GR-LBD* in the presence of Sgt1, Hsp90 and Aha1 by aUC sedimentation is shown. Normalized c(S) distributions were plotted against the sedimentation coefficient S [GR-LBD*: 500 nM, Hsp90: 10 µM, Sgt1: 20 µM, Aha1 10 µM, ATP: 5 mM]. Right panel: Quantification of free GR-LBD was performed by integrating the area under the free GR peak at 2.5 S. The quantification represents means ± SD determined from three independent experiments. E) ATPase analysis of Hsp90 in the presence of Sgt1, Aha1 and GR-LBD [Hsp90: 2 µM, Aha1: 2 µM, Sgt1 8 µM, GR-LBD: 2 µM]. The quantification represents means ± SD determined from three independent experiments. F) Analysis of client complexes in the presence of Sgt1 and Aha1. aUC sedimentation experiment as described in D, but using the Sgt1 truncation Sgt1-TPR-CS. G) ATPase measurement of Hsp90 in the presence of Sgt1-TPR-CS, Aha1 and GR-LBD. [Hsp90: 2 µM, Aha1: 2 µM, Sgt1-TPR-CS: 8 µM, GR-LBD: 2 µM]. The quantification represents means ± SD determined from three independent experiments.

We then investigated how the presence of a client influences this competition using labeled GR-LBD. Upon addition of Hsp90, the majority of GR-LBD is bound to Hsp90, leaving only a small fraction (∼15%) unbound (Figure 5D). The additional presence of Sgt1 leads to the uniform formation of a ternary complex consisting of GR-LBD, Hsp90, and Sgt1, completely replacing the GR-LBD/Hsp90 complex, suggesting a remarkably high affinity of the ternary complex. In contrast, the addition of Aha1 instead of Sgt1 releases the majority of GR-LBD from Hsp90, drastically increasing the proportion of free GR-LBD to nearly 80% (Figure 5D). Previously, it was thought that Aha1 and GR-LBD were not able to bind to Hsp90 at the same time.^23^ However, we can identify a peak shift of the GR-LBD/Hsp90 complex upon Aha1 addition around 1 S, indicating that while GR-LBD is mainly released from Hsp90, a complex of GR, Hsp90, and Aha1 is possible. The addition of Sgt1 to the mixture significantly inhibits the client release by Aha1, reducing it by 35% when both co-chaperones are present in equimolar amounts (Figure 5D). This inhibitory effect is concentration-dependent (Figure S6A, B) and persists in the presence of ATP or ATPγS (Figure 5D, Figure S6C). ATPase activity assays of Hsp90 mirrored these findings. The addition of Aha1 increased the Hsp90 ATPase rate by approximately fourfold. However, this increase is lower in the presence of Sgt1, reflecting the competition of the two co-chaperones for Hsp90 binding (Figure 5E). Notably, the decreased stimulation is substantially more pronounced in the presence of client protein (GR-LBD). To determine whether the protective effect of Sgt1 relies solely on the competition of its CS domain with Aha1 for Hsp90 binding or also involves the SGS domain, we tested a Sgt1 construct containing only the CS and TPR domains. This construct showed a reduced but still significant inhibition of both Aha1-induced client release (Figure 5F) and cycling acceleration (Figure 5G) compared to full-length Sgt1. It remained as effective as full-length Sgt1 in displacing Aha1 from Hsp90 (Figure S6D). These findings suggest that the CS and SGS domains work together to stabilize the Hsp90-client interaction and protect it from dissociation induced by Aha1, effectively prolonging the Hsp90-client interaction.

## DISCUSSION

Despite its essential function in yeast, the role of Sgt1 in the Hsp90 cycle has remained elusive. Our study defines the functional mechanistic of Sgt1 and its critical structural features. Our *in vivo* analysis identified the SGS domain as critical for Sgt1’s essential function. AlphaFold 2 predicts a folded region within the SGS with a relatively high pLDDT (> 80) containing two helices and a constrained C-terminal loop. Our experimental NMR data confirm that the SGS encompasses two helical regions but showed that despite the helical regions, the SGS is conformationally heterogeneous and does not adopt a single conformation. We could pinpoint the essential function of Sgt1 to the second helical motif plus eight subsequent amino acids (up to residue 372) within the SGS domain. Interestingly, this motif is required for Sgt1’s chaperone function, strongly suggesting that the chaperone function underpins its essential role. While the region is dynamic, residue S372 is predicted to create hydrogen bonding with the nearby second helix. High-confidence AlphaFold models of intrinsically disordered regions often predict their conformations in a stabilized state as adopted in the presence of partners, for example.^63^ The AlphaFold model may suggest an SGS conformation when bound to partners. This conformation is likely also present in the apo SGS in low propensity but becomes too sparsely populated in the mutant Sgt1-371, altering its function to a point where it cannot sustain yeast survival.

We found that yeast Sgt1-CS interacts with two distinct binding sites on Hsp90. The known Hsp90-NTD binding site^46^ showed relatively low affinity compared to the new binding site on the Hsp90-MD. The crystal structure of the Hsp90-MD and Sgt1-CS complex revealed a similar interface as observed for the human co-chaperone NudC^53^, but here, the opposite side of the CS beta-sandwich domain interacts with Hsp90. In full-length Hsp90 in the open state, both binding sites are occupied when adding Sgt1. This shows that the binding sites are not mutually exclusive. Several other Hsp90 co-chaperones feature a CS domain, including p23 (Sba1 in yeast) and NudC in humans. However, these lack the SGS domain. Nevertheless, a similar overall architecture is found, where the CS domain is followed by amphipathic helices within a largely disordered region. In these proteins, the CS domain mediates Hsp90 interaction, while the helices in the unstructured region are involved in interactions with clients and Hsp90.^53,54,56^ For Sgt1, we found that the SGS domain is involved in client binding and in transient interaction with the Hsp90-CTD. This interaction, together with the client binding properties of the SGS, seems to be sufficient to support viability at high expression levels. However, the CS and SGS domains are essential for viability at native expression levels. This suggests that the CS-mediated interaction with Hsp90 is necessary for the SGS domain to execute its essential function. An additional client binding function of Sgt1 has been assigned to the Sgt1-TPR domain, which binds Skp1, a component of the kinetochore and SCF complex.^26,30,40,64^ Although Skp1 is essential in yeast, our findings suggest that its TPR interaction with Sgt1 is not essential for viability, in contrast to previous findings.^64^ However, they align with the fact that the TPR domain is the least conserved domain and completely absent in several species.^44^ Instead of the TPR domain, our data suggest that the interaction of the SGS domain with client proteins is essential for its function.

Sgt1 neither affects the ATPase activity of Hsp90, nor does it exhibit a pronounced interaction with Hsp70 or JDPs, indicating that it is not involved in client transfer from the Hsp70 system. Although we found that Sgt1 can directly interact *via* its SGS domain with the client GR, we did not identify evidence that it promotes the binding of GR to Hsp90. This speaks against a client loading function. However, we see that client processing towards the native, hormone-binding conformation becomes more efficient in the presence of Sgt1. Since client transfer and loading are not affected, it is reasonable to assume that client binding by Sgt1 is involved in the later steps of the Hsp90 cycle. In line with this concept, we detected a stabilization of Hsp90-GR complexes in the presence of Sgt1.

Previously, we found that the co-chaperone Aha1 competes with GR for binding to Hsp90.^23^ Deletion of Aha1 in yeast results in increased levels of functional GR.^18^ This also holds true for other proteins.^18–20,22^ Similarly, the absence of Aha1 in mammalian cells improves the folding efficiency and stability of both wild-type and disease-associated CFTR.^15,21^ Our results show that Sgt1 counteracts this process by effectively impairing Aha1 binding and inhibiting Aha1-induced client release. The superimposition of our structure of the Hsp90-MD/Sgt1-CS complex over the previously reported Hsp90/Aha1 structures^16^ demonstrates that the two co-chaperones cannot bind simultaneously. Indeed, the CS domain of Sgt1 clashes with the binding of the Aha1-NTD to Hsp90 in both the open and closed conformation (Figure S6E). Our results thus suggest that the two co-chaperones target later steps in the Hsp90 chaperone cycle but with opposing effects. Supporting this notion, overexpression of Aha1 in a temperature-sensitive Sgt1 mutant led to a substantial growth defect in yeast cells. This effect was abolished when an Aha1 mutant insufficient to bind to Hsp90 was used.^65^ Thus, Sgt1 appears to keep the client release effect of Aha1 at bay. In the absence of Sgt1, Aha1 will reduce the dwell time of GR. Thus, Sgt1 stabilizes the Hsp90-client complex and extends the processing phase of Hsp90 for clients by shielding them from Aha1 (Figure 6).

**Figure 6.**
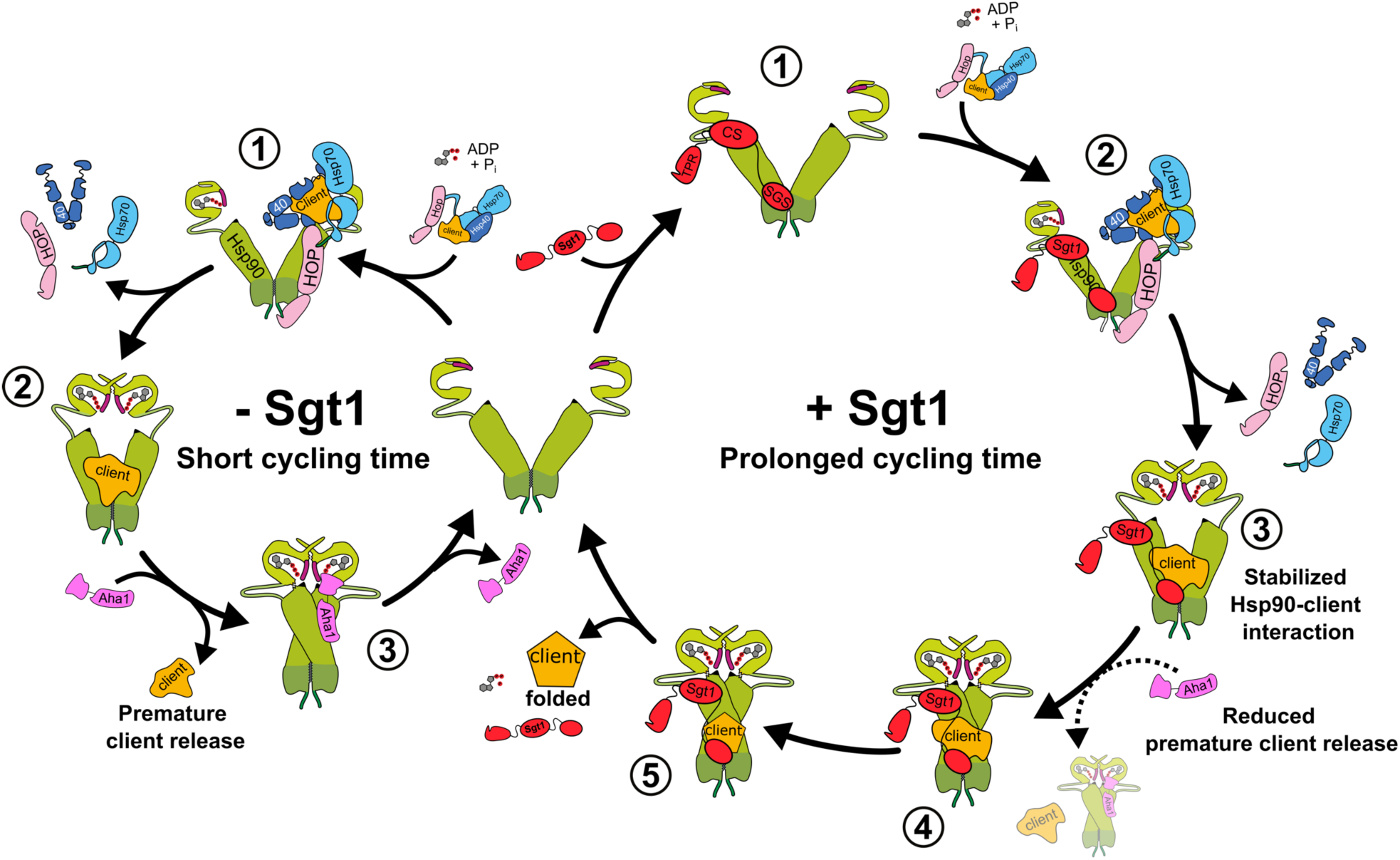
Schematic model of how Sgt1 influences the client maturation process. A schematic representation of Sgt1’s role in the yeast Hsp90 cycle is shown. For clarity, yeast Sgt1 is depicted as a monomer, although it is capable of forming dimers. Left panel: In the absence of Sgt1, client proteins are transferred from the Hsp40/Hsp70 complex to Hsp90 with the help of Hop (steps 1–2). The co-chaperone Aha1 can bind to Hsp90 without restriction, accelerating the Hsp90 cycling time and promoting premature release of clients (step 3). Right panel: In the presence of Sgt1, client proteins are loaded onto Hsp90 via Hop (steps 1–2), as in the canonical cycle. However, Sgt1 stabilizes the Hsp90-client interaction and prevents excessive Aha1 binding (step 3). This slows the Hsp90 cycling time as indicated by a larger cycle and increases the dwell time of clients in the Hsp90 complex. Thus, Sgt1 promotes the proper maturation of the client protein (steps 4-5).

### Limitations of the study

While our study provides important insights into the structural and functional role of Sgt1, several limitations remain. First, we focused exclusively on yeast Sgt1, and further studies are needed to determine whether homologs in other organisms, such as humans, function in a similar manner. Second, a more detailed structural analysis of the Sgt1 interaction with the client GR-LBD, as well as the ternary complex with Hsp90, could provide deeper insights into its structure-function relationship. Finally, whether there is a specificity of Sgt1 for certain client proteins remains to be determined.

## ACKNOWLEDGMENTS

We thank Kaya Bergmann for laboratory support, and we are particularly grateful to Sam Asami and Gerd Gemmecker (TUM) for support with NMR experiments. We acknowledge access to NMR measurement time at the Bavarian NMR Center. This study was supported by the German Research Foundation DFG (SFB1035, Projektnummer 201302640, project A03). S.E. acknowledges a doctoral fellowship from the Studienstiftung des deutschen Volkes.

## AUTHOR CONTRIBUTIONS

Conceptualization, F.D., S.E., C.D., A.L., J.B., and M.S.; Methodology, S.E., F.D., C.D., A.L.; Investigation, F.D., S.E., C.D., A.L., V.N., G.M.P. and O.F.; Visualization, S.E., F.D., and O.F.; Writing – Original Draft, S.E. and F.D.; Writing – Review & Editing, J.B., R.R., M.S. and O.F.; Supervision, J.B., M.S., and R.R.; Resources and Funding Acquisition, J.B., M.S., and R.R.

## STAR METHODS

### RESOURCE AVAILABILITY

#### Lead contact

Further information and requests for resources and reagents should be directed to, and will be fulfilled by, the lead contact, Michael Sattler and Johannes Buchner.

#### Materials availability

Plasmids generated in this study will be made available on request, but we may require a payment and/or a completed Materials Transfer Agreement if there is potential for commercial application.

#### Data and code availability

- All data and any additional information required to reanalyze the data reported are available from the corresponding authors and/or are included in the manuscript.
- The crystal structure of the Hsp82-MD/Sgt1-CS complex have been deposited to the Protein Data Bank with the accession number PDB: 9Q8O

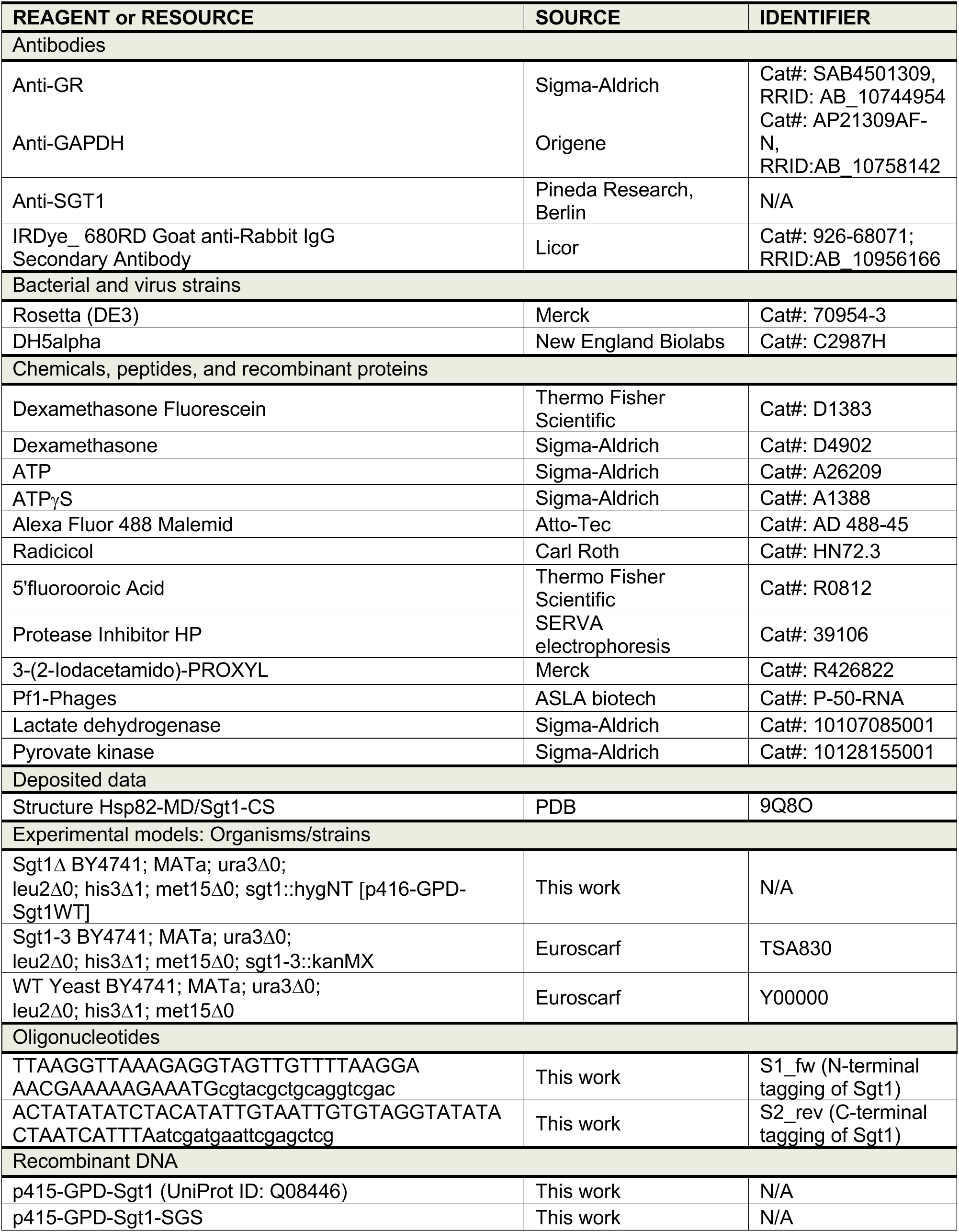
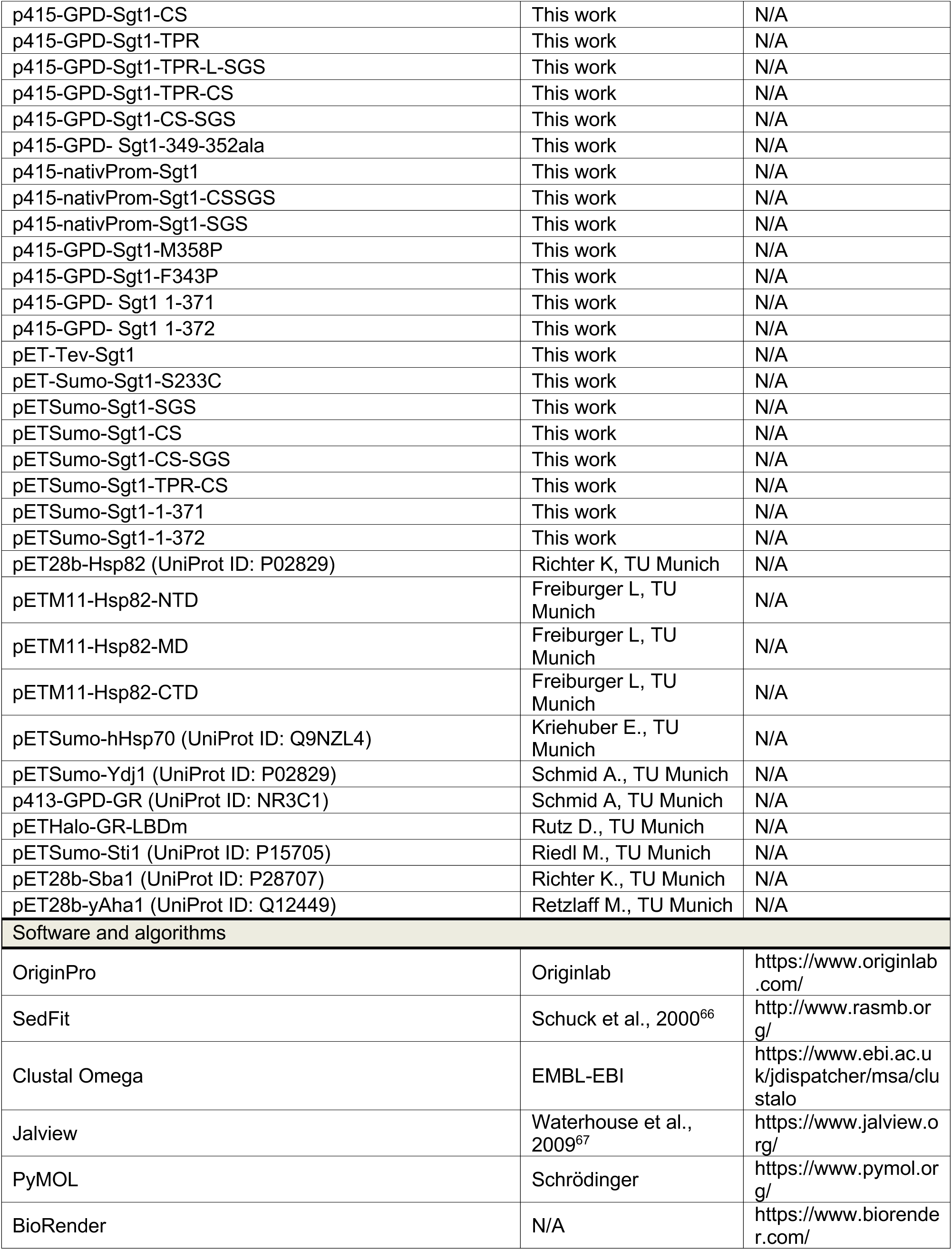
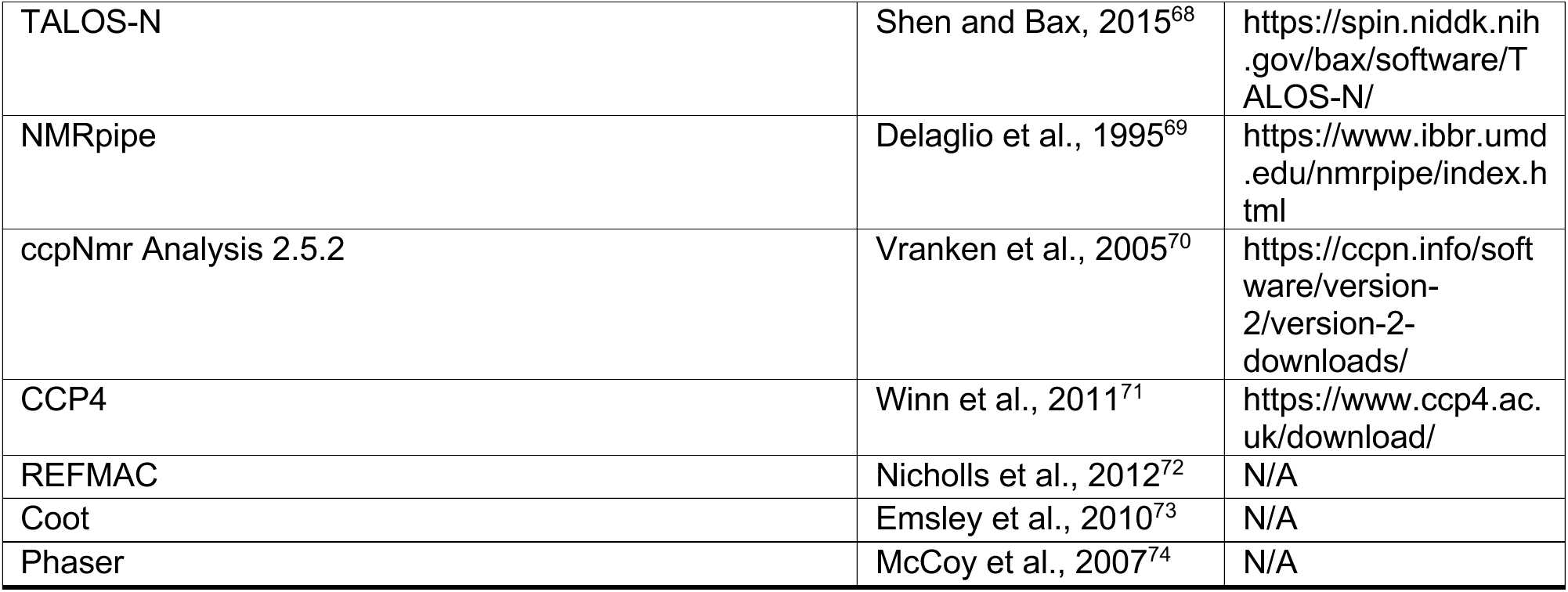
KEY RESOURCES TABLE.

## EXPERIMENTAL MODEL AND SUBJECT DETAILS

### Yeast strains

All yeast strains used in this study are derivatives of BY4741 (MATa; ura3δ0; leu2δ0; his3δ1; met15δ0) and with the exception of the Sgt1δ strain, all strains were obtained from Euroscarf. Chromosomal deletion of Sgt1 was performed using a PCR-based method as described by Janke et al. 2004.^75^ Specifically, the primers S1 and S2 were designed to replace the Sgt1 gene with the hphNT1 marker from the pFA6a-hphNT1 plasmid. The PCR product was used for yeast transformation, and colonies positive for the hphNT1 marker were confirmed by colony PCR.

## METHOD DETAILS

### Yeast transformation

Yeast strain transformation was performed as described by Gietz and Schiestl 2007^76^, in a slightly modified way. In brief, yeast cells were grown over night in YPD medium and re-inoculated in 25 mL to a starting OD_600_=0.15. Yeast growth continued for approximately two cell division cycles at 30°C (temperature-sensitive mutants were grown at 25°C). Afterwards, cells were harvested (3000 x g for 5 min), washed once with sterile water, subsequently washed with 1 mL 0.1 M lithium acetate and then resuspended in 0.5 mL 0.1 M lithium acetate. 50 µL of the cell suspension was used for a single transformation reaction. 240 μL PEG-3350, 36 μL of 1 M lithium acetate, 10 μL ssDNA and 72 – X μL H2O were added with X representing the volume of DNA solution. After mixing, the sample was incubated at 30°C for 30 min and at 42°C for 30 min. Cells were then harvested at 4000 x g for 30 s and plated on selective medium.

### Plasmid shuffling

The well describe method of 5-Fluoroorotic acid (5-FOA) plasmid shuffling was used to characterize essential genes.^77^ The *sgt1Δ* [p416-GPD-SGT1] strain was used for plasmid shuffling, in which the genomic copy of SGT1 was deleted and replaced by a URA3-marked plasmid carrying a wild-type SGT1 gene (p416-GPD-SGT1). The URA3 gene encodes orotidine-5’-phosphate decarboxylase which converts 5-FOA to the cell toxic 5-fluorouracil allowing counter-selection. The strain was transformed with a second plasmid containing Sgt1 mutants and plated on selective medium without 5-FOA. Appearing colonies were streaked out on plates containing 0.1% w/v 5-FOA to select for loss of the p416-GPD-SGT1 plasmid.

### Protein expression, purification and labeling

<u>Sgt1, Sgt1 mutants and domains, Hsp82 domains, Sti1, hHsp70 and Ydj1</u> were expressed with a 6xHis-SUMO tag or a 6xHis-TEV tag in *E. coli* Rosetta cells. Cells were lysed in NiNTA buffer A (50 mM Na2HPO4 pH 7.5, 300-500 mM NaCl, 10 mM imidazole, 1 mM DTT + protease inhibitor HP, 2 mM PMSF). Lysates were cleared at 40,000 g for 1 h. Cleared lysates were subjected to affinity chromatography on a Hi-Trap FF column. Bound protein was washed with 2 mM ATP dissolved in NiNTA buffer A before elution with NiNTA buffer B containing 300 mM imidazole. The elute fractions were pooled and the SUMO tag was cleaved by addition of Ulp1 or Tev protease (depending on the construct) and dialysis in NiNTA A buffer (10 mM imidazol) over night at 4°C. To get rid of the SUMO-/His-tag and the protease, the digested proteinwas run again over a His-Trap FF column and the flow through was collected. Hsp82 and full length Sgt1 were diluted with Resource Q buffer A (40 mM HEPES, pH 7.5, 20 mM KCl, 1 mM EDTA, 1 mM DTT) to a salt concentration lower than 50 mM and loaded on to a Resource Q ion-exchange column. Elution of the bound proteins was done using a continuous salt gradient of Resource Q buffer B (40 mM HEPES, pH 7.5, 1 M KCl, 1 mM EDTA, 1 mM DTT). Subsequently all proteins were subjected to size exclusion chromatography in SEC buffer (40 mM Hepes, pH 7.5, 150 mM KCl, 5 mM MgCl2, 1 mM DTT). <u>Hsp82, Sba1 and</u> <u>Aha1</u> were expressed with a C-terminal 6xHis-tag. Affinity chromatography was performed as described above. Subsequently the proteins were directly subject to ion-exchange and size exclusion chromatography. All experiments with <u>GR-LBD</u> were conducted with the stabilized, purified GR-LBDm (aa 527-777, F602S/A605V/V702A/E705G/M752T), purified as previously published.^78^

Proteins were fluorescently labeled with maleimide-Atto488. Proteins were dialyzed into a non-reducing reaction buffer (40 mM Hepes, pH 7.5, 150 mM KCl, 5 mM MgCl2). The dye was added to a molar ratio of protein:dye (1:2) and incubated for 1-2 h at RT. The reaction was quenched by adding 5 mM DTT. Free dye was separated by a Sephadex® column (10 mm x 300 mm) using SEC buffer (40 mM Hepes, pH 7.5, 150 mM KCl, 5 mM MgCl2, 1 mM DTT).

### Analytical Ultracentrifugation

Analytical ultracentrifugation experiments were performed using a ProteomLab Beckman XL-A centrifuge (Beckman Coulter, Brea, California) equipped with an AVIV fluorescence detection system (Aviv biomedical Inc., Lakewood, USA). Experiments were performed at 42,000 rpm and 20°C in an eight-hole Ti-50 Beckman-Coulter rotor. 300 scans were recorded, in a total measurement time of 6h. If not stated otherwise, 500 nM Atto488 labeled protein was detected in aUC buffer (20 mM Hepes pH 7.5, 20 mM KCL, 5 mM MgCl2). All other protein concentrations were used as indicated in the figure legends. Nucleotides were added to a concentration of 5 mM (ATP) and 2 mM (ATPγS). Sedfit and OriginPro 2024 were used for data analysis. The data was normalized to the total area under the curve whereby, peaks of free label were excluded from the calculation. To quantify the percentage of specific complexes, normalized data was used and the area under the peak of the desired complex was determined.

To determine the dissociation constant (K_D_) of Sgt1 and Hsp82, the percentage of bound species, was quantified by calculating the area under the Hsp82-Sgt1 complex. The concentration of Hsp82 was plotted against the percentage of bound species. The Hill1 fit was used to determine the dissociation constant (KD) and the hill coefficient (N). Mean±Standard deviation was determined by three individual experiments.

### Fluorescence Polarization

Hormone binding to the GR-LBD was monitored as described previously.^52^ Briefly, for equilibrium GR-LBD measurements, 1 µM of apo GR-LBD was mixed with different combinations of the following protein: 2 µM Hsp40 (Ydj1), 10 µM hHsp70, 4 µM Hsp90 (Hsp82), 12 µM Sgt1 and 5 mM ATP in 40 mM Hepes pH 7.5, 150 mM KCl, 5 mM MgCl2, 2 mM DTT buffer. The proteins were incubated at room temperature for 60 min. F-Dex was added in a final concentration of 100 nM and the binding of F-Dex to the GR-LBD was monitored at 30°C by measuring the fluorescence polarization values. For the analysis, the start-value immediately after addition of F-Dex was subtracted from the trace. The last 10 values were used to calculate the end points of each measurement.

### ATPase assay

The Hsp82 ATPase activity was measured with a regenerative ATPase assay as previously published.^79,80^ Briefly, assay buffer was prepared containing 5.17 mM phosphoenolpyruvate (PEP), 0.43 mM nicotinamidadenine-dinucleotidephosphate (NADH), 5.17 U/mL pyruvate kinase (PK) and 26.06 U/mL lactate dehydrogenase (LDH) in 40 mM Hepes, pH 7.5, 45 mM KCl, 5 mM MgCl2. Assay buffer was mixed 1:1 with proteins diluted in 40 mM Hepes, pH 7.5, 45 mM KCl, 5 mM MgCl2 and incubated for 30 min at 30°C. Concentrations of the proteins was used as indicated. The reaction was initiated by the addition of 2.5 mM ATP and the absorption at 340 nm was continuously recorded at 30°C. As a control, each protein batch was tested for ATPase activity to exclude impurities that cause an intrinsic ATPase activity of the proteins. Hsp82 was tested by addition of 50 µM Radicicol, and the background activity was subtracted from the measurement. Three independent experiments were performed.

### Western Blot

For western blot 5 OD unites of an exponentially growing yeast cultures were lysed according to the alkali lysis method. The pellet fraction was solubilized in Laemmli buffer. A total of 40 µg of protein was loaded on a SDS gel and blotted on a PVDF membrane. Peroxidase coupled secondary antibodies were used and bands were detected using ImageQuant LAS4000. Three independent experiments were performed.

### NMR

All NMR experiments were performed on Bruker Avance spectrometers equipped with cryogenically cooled TCI probe heads, operating at magnetic field strengths corresponding to ^1^H Larmor frequencies of 600 to 1200 MHz. The sample temperature was set at 298 K for all experiments unless otherwise specified. ^1^H,^15^N correlation experiments were performed in 40 mM phosphate buffer, 100 mM NaCl, 2 mM DTT, pH 6.8, 8% D_2_O, while methyl-labeled experiments were all performed in 20 mM tris-D_11_, 100 mM NaCl, 5 mM MgCl_2_, 2 mM DTT-D_10_, 0.02% NaN_3_, pH 7, in 99.9% D_2_O. All data were processed with NMRpipe^69^ and analyzed in CCPNMR^70^.

### Chemical shift assignments

The backbone assignment of all Sgt1 constructs was performed using standard 3D heteronuclear experiments HNCACB, HNCA, CBCA(CO)NH or (H)C(CCO)NH, HNCO and HN(CA)CO.^81^ The combined ^13^Ca, ^13^Cb, ^13^C’ secondary shifts (Δδ_av_(^13^C)) were calculated as [Δδ(^13^Cα) – Δδ(^13^Cδ) + Δδ(^13^C’)], where Δδ is the difference between observed ^13^C chemical shifts and random coil values. Since Δδ(^13^Cα) and Δδ(’^13^C’) are positive and Δδ(^13^Cδ) is negative in helical conformations and opposite in strand conformations, the value Δδ_av_(^13^C) can be used to evaluate secondary structure propensity.^82^ Methyl-labeled samples were prepared as previously described. Backbone and methyl resonances of Hsp82 were previously assigned by us.^23,83^ The {^1^H}–^15^N heteronuclear NOE were recorded as described ^84^ in an interleaved manner with a recycling time of 4s.

### Paramagnetic relaxation enhancements (PRE)

The mutant Sgt1-SGS E375C was first exchanged to an NMR-buffer without DTT using a PD10 buffer exchange column (GE Healthcare, Buckinghamshire, UK). Two other mutants were generated (344C and 366C) but were not included as they significantly affected the ^1^H-^15^N HSQC spectra of SGT1-SGS and thus likely affected its conformational state. IPSL (N-(1-oxyl-2,2,5,5-tetramethyl-3-pyrrolidinyl)iodoacetamide) was added to a 10 times molar excess to Sgt1-SGS E375C in non-reduced buffer and incubated overnight at room temperature and protected from light. The excess spin label was removed using a PD10 column. 200 μM of Sgt1-SGS E375C IPSL-labeled was prepared and measured. Data were recorded using ^1^H,^15^N HSQC experiment with a recycling delay of 4s. 10 times molar excess of fresh ascorbic acid was added to reduce the paramagnetic probe, and the experiment was repeated using the same parameters to record the reference experiment in the absence of PRE.^85,86^ Theoretical PRE effects on Sgt1-SGS E375C were calculated based on the AF2 model from the AF-PSD (AF-Q08446-F1-v4). Seven different conformations of IPSL covalently bound to C375 were generated using the python package chiLife.^87^ For each structure and conformation, the distance between the paramagnetic center and the backbone NH of Sgt1-SGS S375C were extracted in pymol using the distancetoatom script. The relaxation ratio was calculated as described^85^ for each conformation, assuming a tumbling correlation τ_c_ of 50 ns, an ^1^H-R_2_ transverse relaxation rate of 80 s^-1^, and a total INEPT evolution time of 9 ms.

### Residual dipolar couplings

^1^H-^15^N residual dipolar couplings (RDCs) were obtained by recording IPAP-HSQC experiments on ^15^N-labeled samples of Sgt1-SGS at 270 µM concentration, in absence and in presence of 12.5 mg/mL of Pf1-phages (ASLA biotech). The alignment was confirmed by ^2^H 1D NMR that showed a deuterium splitting value of around 9 Hz.

### Titrations

Titration of the ^15^N-labeled Sgt1-CS-SGS were performed at 100 µM with either 2-points titration 1:0.5 and 1:1 for Hsp90-FL, Hsp90-CTD and GR or 7-point titrations for Hsp90-NTD and Hsp90-Md. Reported CSPs on the figure correspond to the 1:1 ratios. Perturbations and intensity changes on methyl-labeled Hsp90 were performed at 300 µM with a single addition of Sgt1 constructs at 450 µM. Titrations were performed with proteins purified in identical buffers. The assignment of fully bound-forms, used to determine chemical shifts perturbations, was performed from the successive addition of binding partners and following the chemical shift perturbations. All chemical shift perturbations were calculated as a weighted average following the equations CSP = ((Δδ^1^H)^2^ + (Δδ^15^N^∗^0.15)^2^)^1/2^ for ^1^H, ^15^N spectra and CSP = ((Δδ^1^H)^2^ + (Δδ^13^C^∗^0.3)^2^)^1/2^ for ^1^H, ^13^C spectra.^88^

### Crystallography

Initial screening of crystallization conditions was carried out by the vapor diffusion method. Sitting drops were set up using 400 nL of a 1:1 mixture of protein and crystallization solutions. The co-crystallization was performed using Hsp90-MD and Sgt1-CS with a complex concentration of 20 mg/mL. Crystals were obtained at 20°C from a solution containing 0.04 M Magnesium chloride, 0.05 M Sodium cacodylate, 5% v/v 2-Methyl-2,4-pentanediol. Single crystals were cryoprotected using 25% glycerol in the mother solution and flash-frozen in liquid nitrogen. The diffraction data were collected at the X06DA beamline at the Swiss Light Source (Paul Scherrer Institut, Villigen, Switzerland). The data were indexed and integrated using the XDS program package^89^ and scaled and merged using the Aimless program^90^ The initial phases were obtained by molecular replacement, calculated using Phaser software^74^ using Hsp90-MD structure (pdb:1HK7^24^) and the Sgt1-CS from Arabidopsis thaliana (pdb:2JKI^46^). All crystallographic calculations were carried within the CCP4 program suite.^71^ The protein structure was refined with the program REFMAC^72^ and manual adjustments were made to the models using Coot.^73^ Atomic coordinates and structure factors for the Hsp90-MD/Sgt1-CS complex have been deposited in the RCSB Protein Data Bank with pdb code 9Q8O.

