## Supplementary_Figures_S1-S6 for "The essential co-chaperone Sgt1 regulates client dwell time in the Hsp90 chaperone cycle"

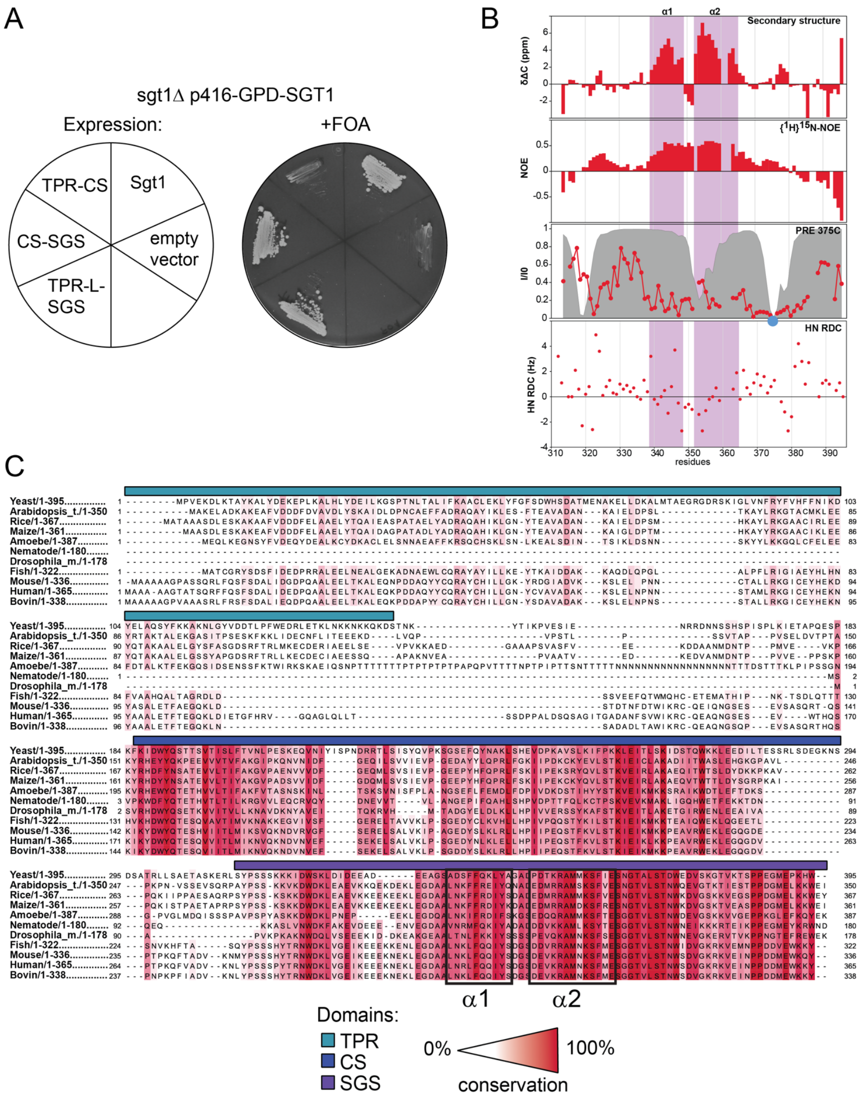


**Figure S1. The SGS lacks a defined tertiary conformation but contains two highly conserved helices, related to Figure 1.**

A) Plasmid shuffling (right panel) of the BY4741 sgt1Δ p416-GPD-SGT1 strain transformed with p415-GPD plasmids containing Sgt1, Sgt1 domain constructs or empty vector as indicated in the left panel.

B) Data from Figure 1C with the addition of RDCs along the sequence of Sgt1-SGS with alignment from Pf1-phages. Helices location is shown in pink.

C) Multiple sequence alignment (MSA) generated by Clustal Omega (https://www.ebi.ac.uk/jdispatcher/msa/clustalo). The amino acid sequence of the Sgt1 homologs of Yeast (UniProt: A6ZNQ9), Arabidopsis thaliana (UniProt: Q9SUR9), Rice (UniProt: Q0JL44), Maize (UniProt: A0A3L6DPG1), Amoebe (Dictyostelium discoideum; UniProt: Q55ED0), Nematode (Parascaris univalens; UniProt: A0A915BFG5), Dorsophila melanogaster (UniProt: Q8SY87), Fish (Danio rerio, UniProt: Q5U3E4), Mouse (UniProt: Q9CX34), Human (UniProt: Q9Y2Z0), Bovin (UniProt: Q2KIK0) were used for the analysis. The color intensity represents the conservation of the residues across the species.

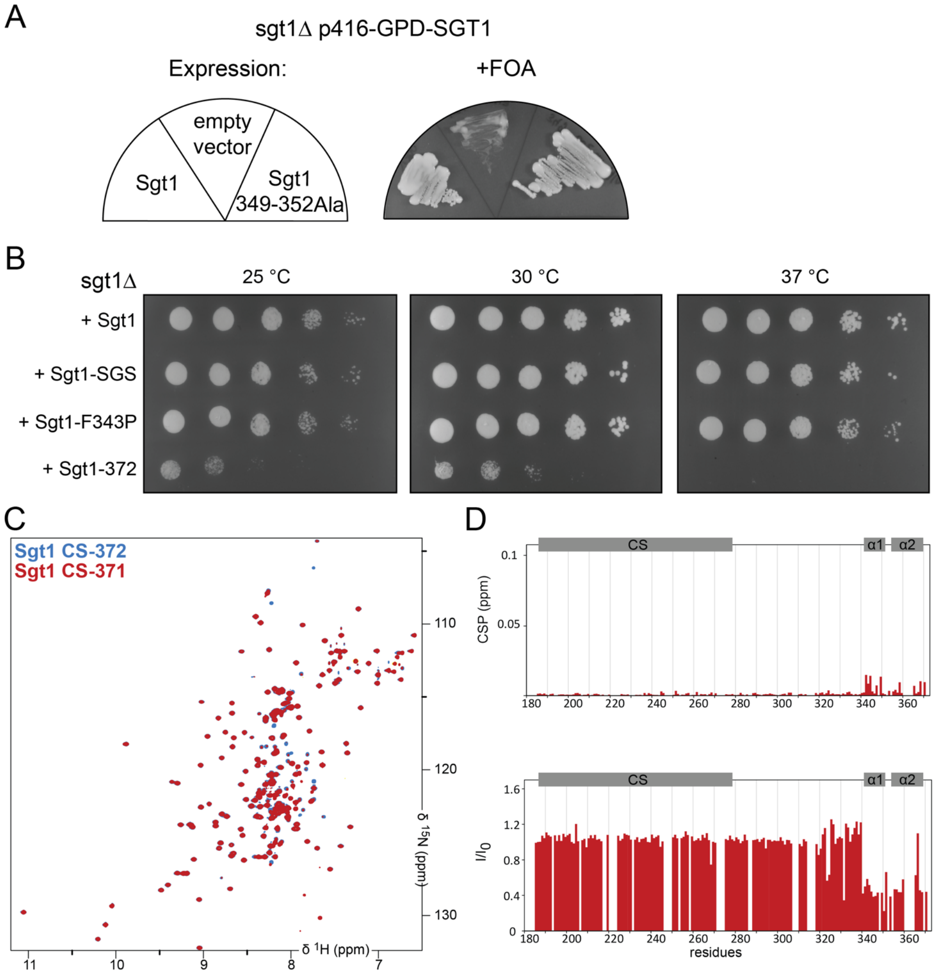


**Figure S2. Residue 372 stabilizes the helical conformations of the SGS domain, related to Figure 2.**

A) Plasmid shuffling (right panel) of the BY4741 sgt1Δ p416-GPD-SGT1 strain transformed with p415-GPD plasmids containing Sgt1, Sgt1 mutant or empty vector as indicated in the left panel.

B) Drop assay of viable Sgt1 shuffling strains. Sgt1Δ yeast strains expressing Sgt1, Sgt1-SGS, Sgt1-F343P, Sgt1-372 on the plasmid p415-GPD were generated by plasmid shuffling. The shuffled strains were grown in liquid SD media at 30°C and dropped on SD agar in 10-fold serial dilution. Plates were incubated 2 days at the indicated temperatures.

C) $^{1}H$-$^{15}N$ HSQCs overlays of CS-372 (blue) and CS-371 (red) showing the differences induced by the early truncations.

D) CSPs and intensity ratios from $^{1}H$-$^{15}N$ HSQC spectra of CS-372 vs CS-371 (red). CS-371 shows lower intensity in the helical region, which is likely due to a change in dynamics and an increased intermediary exchange contribution.

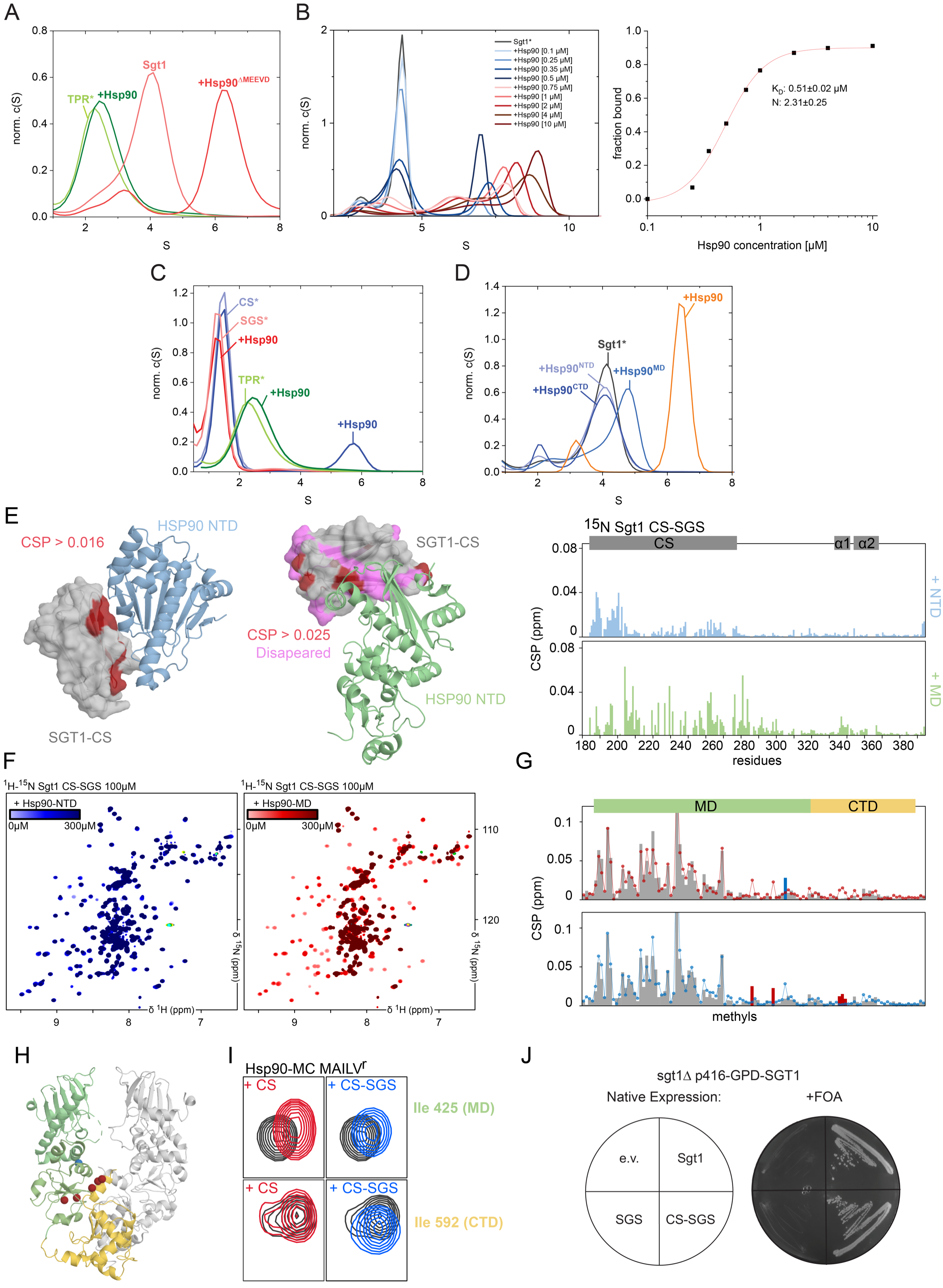


**Figure S3. Characterization of the interaction between Sgt1 and Hsp90, related to Figure 3.**

A) Complex formation analysis by aUC sedimentation of the labeled TPR domain of Sgt1 in presence of Hsp90 or labeled Sgt1 in presence of an $Hsp90^{\Delta MEEVD}$ mutant. Normalized c(S) distributions were plotted against the sedimentation coefficient S [TPR*: 500 nM, Hsp90: 5 µM, Sgt1*: 500 nM, $Hsp90^{\Delta MEEVD}$: 5 µM, ATP: 5 mM].

B) Determination of the dissociation constant between full length Sgt1 and Hsp90. Left panel: Representative complex formation analysis of labeled Sgt1* and varying concentrations of Hsp90 by aUC sedimentation. Normalized c(S) distributions were plotted against the sedimentation coefficient S [Sgt1*: 500 nM, Hsp90: concentrations as indicated, ATP: 5 mM]. Right panel: Quantification of the bound fraction was calculated by integration of the area under the Sgt1-Hsp90 complex. Mean ± SD were determined from three independent experiments.
C) Complex formation analysis of labeled Sgt1 domains (SGS*, CS*, TPR*) in the presence of Hsp90 by aUC sedimentation. Normalized c(S) distributions were plotted against the sedimentation coefficient S [SGS*/CS*/TPR*: 500 nM, Hsp90: 5 µM, ATP: 5 mM].
D) Complex formation analysis of labeled Sgt1* in the presence of Hsp90 full length and domains by aUC sedimentation. Normalized c(S) distributions were plotted against the sedimentation coefficient S [Sgt1*: 500 nM, Hsp90: 5 µM, Hsp90-NTD/-MD/-CTD: 5 µM, ATP: 5 mM].
E) CSP plots of resonances from the $^{1}H$-$^{15}N$ HSQC of the equimolar addition to Sgt1-CS-SGS of Hsp90-NTD (blue), Hsp90-MD (green) mapped onto the structure model of the Hsp90-NTD/Sgt1-CS derived from the 2JKI pdb structure and from the Hsp90-MD/Sgt1-CS structure reported here (Fig. 2B).
F) Spectra showing the full titration of Sgt1-CS-SGS with Hsp90-NTD (blue) and Hsp90-MD (red) showing the stronger affinity of Hsp90-MD.
G) Methyl CSP from the titration to Hsp90-MC of Sgt1-CS (top panel, bars) and Sgt1-CS-SGS (bottom panel bars). The line plots report CSPs from the other for ease of comparison (Sgt1-CS-SGS in red and Sgt1-CS in blue). The strongly different CSPs between the two titrations are shown in red (higher in Sgt1-CS-SGS) or blue (higher in Sgt1-CS).
H) Report of the CSP from G) on the structure of Hsp90-MC (from 2cg9) using the same colors.
I) Extract of the $^{1}H$-$^{13}C$ methyl spectrum of Hsp90-MC with either Sgt1-CS or Sgt1-CS-SGS shows additional perturbations in the CTD induced by the presence of the SGS region.
J) Plasmid shuffling (right panel) of the BY4741 sgt1∆ p416-GPD-SGT1 strain transformed with the p415 plasmids containing Sgt1, Sgt1 mutant or empty vector under the control of the native Sgt1 promotor as indicated in the left panel.

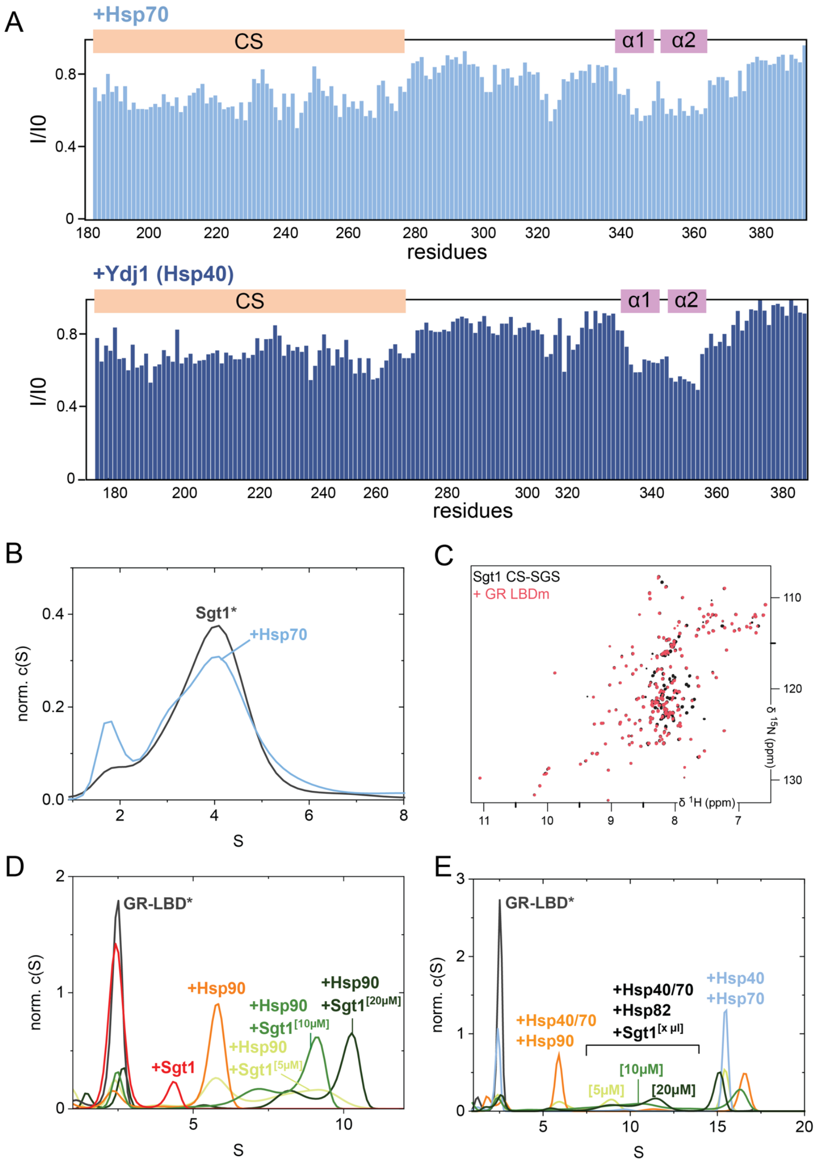


**Figure S4. Interactions of the SGS domain with the Hsp70 system and client proteins, related to Figure 4.**

A) Intensity ratio comparison of resonances from the $^{1}H$-$^{15}N$ HSQC of Sgt1-CS-SGS before and after addition of Hsp70 (top), or Ydj1 (bottom), showing very little interaction.

B) Complex formation analysis of labeled Sgt1* in the presence of Hsp70 by aUC sedimentation. Normalized c(S) distributions were plotted against the sedimentation coefficient S [Sgt1*: 500 nM, Hsp70: 10 µM, ATP: 5 mM].

C) $^{1}H$-$^{15}N$ HSQC spectra of Sgt1-CS-SGS (black) before and after the addition of GR-LBDm (red).

D) Complex formation analysis of the labeled GR-LBD* in the presence of Hsp90 and Sgt1 by aUC sedimentation. Normalized c(S) distributions were plotted against the sedimentation coefficient S [GR-LBD*: 500 nM, Sgt1: as indicated, Hsp90: 10 µM ATP: 5 mM].

E) Complex formation analysis of the labeled GR-LBD* in the presence of Hsp90, Sgt1, Hsp40 and Hsp70 by aUC sedimentation. Normalized c(S) distributions were plotted against the sedimentation coefficient S [GR-LBD*: 500 nM, Sgt1: as indicated, Hsp90: 10 µM, Hsp70: 10 µM, Hsp40: 2 µM, ATP: 5 mM].

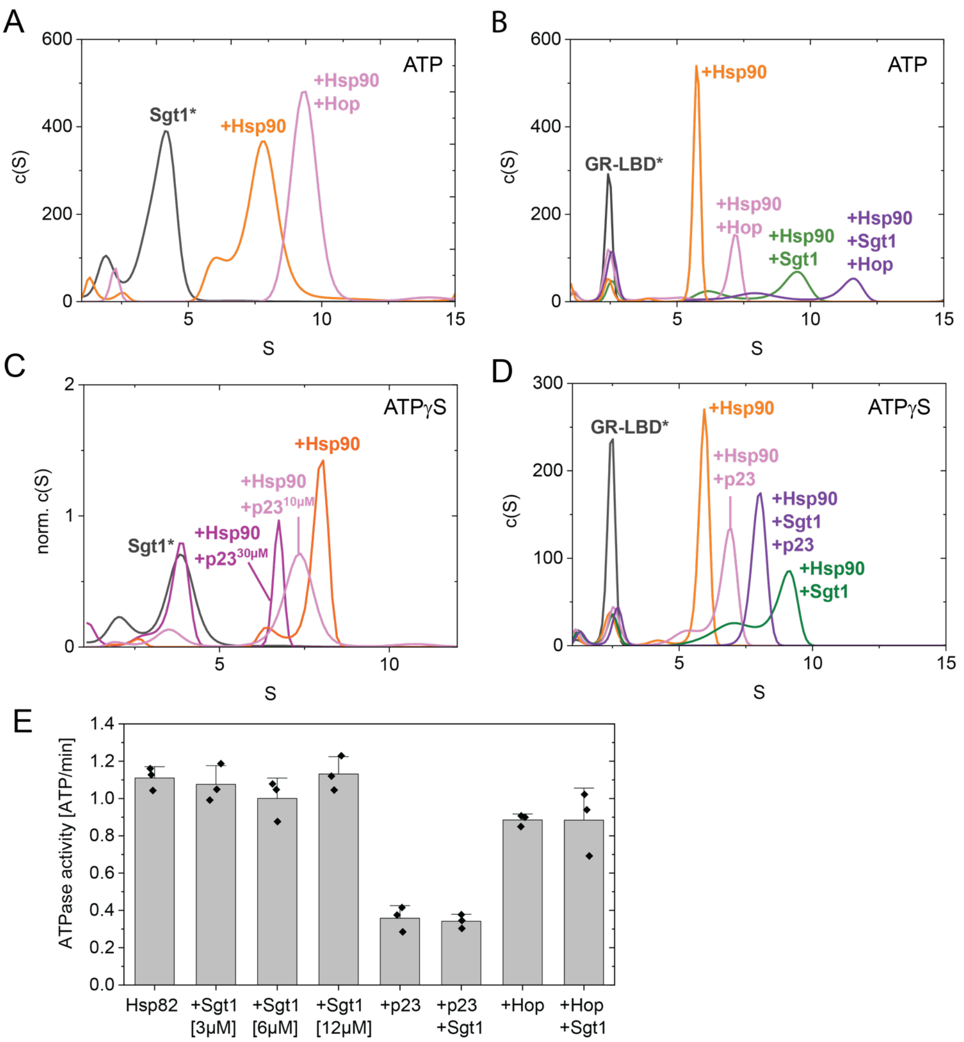


**Figure S5. Interplay of Sgt1 with the co-chaperones Hop and p23, related to Figure 5.**

A) Complex formation analysis of the labeled Sgt1* in the presence of Hsp90, and Hop by aUC sedimentation is shown. c(S) distributions were plotted against the sedimentation coefficient S [Sgt1*: 500 nM, Hsp90: 5 µM, Hop: 10 µM, ATP: 5 mM].

B) Complex formation analysis of the labeled GR-LBD* in the presence of Hsp90, Sgt1 and Hop by aUC sedimentation is shown. c(S) distributions were plotted against the sedimentation coefficient S [GR-LBD*: 500 nM, Hsp90: 10 µM, Hop: 10 µM, Sgt1: 10 µM, ATP: 5 mM].

C) Complex formation analysis of the labeled Sgt1* in the presence of Hsp90, and p23 by aUC sedimentation is shown. Normalized c(S) distributions were plotted against the sedimentation coefficient S [Sgt1*: 500 nM, Hsp90: 5 µM, p23: concentration as indicated, ATPγS: 2 mM].

D) Complex formation analysis of the labeled GR-LBD* in the presence of Hsp90, Sgt1 and p23 by aUC sedimentation is shown. c(S) distributions were plotted against the sedimentation coefficient S [GR-LBD*: 500 nM, Hsp90: 10 µM, p23: 10 µM, Sgt1: 10 µM, ATPγS: 2 mM].

E) ATPase analysis of Hsp90 in the presence of Sgt1, Sba1 and Sti1. [Hsp90: 2 µM, Sgt1: as indicated or 12 µM, Sba1: 2 µM, p23: 2 µM]. The quantification represents means ± SD determined from three independent experiments.

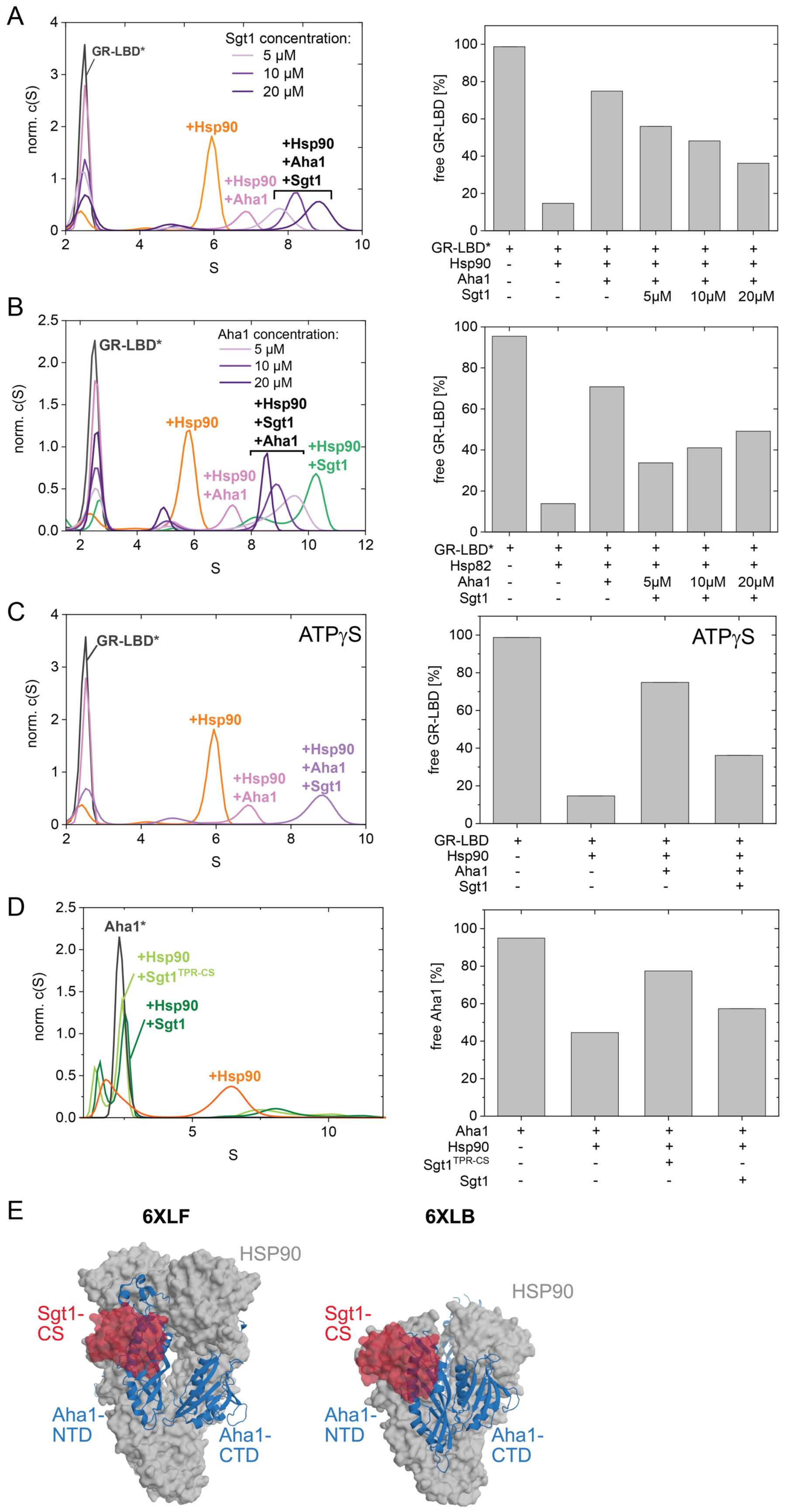


**Figure S6. Characterization of the antagonistic interplay between Sgt1 and Aha1, related to Figure 5.**

A) Complex formation analysis of labeled GR-LBD* in the presence of Aha1, Hsp90 and varying concentrations of Sgt1 by aUC sedimentation is shown. Normalized c(S) distributions were plotted against the sedimentation coefficient S [GR-LBD*: 500 nM, Hsp90: 10 μM, Aha1: 10 μM, Sgt1 concentration as indicated, ATP: 5 mM]. Right panel: Quantification of free GR-LBD was performed by integrating the area under the free GR peak at 2.5 S.
B) Complex formation analysis of labeled GR-LBD* in the presence of Sgt1, Hsp90 and varying concentrations of Aha1 by aUC sedimentation is shown. Normalized c(S) distributions were plotted against the sedimentation coefficient S [GR-LBD*: 500 nM, Hsp90: 10 μM, Sgt1: 20 μM, Aha1 concentration as indicated, ATP: 5 mM]. Right panel: Quantification of free GR-LBD was performed by integrating the area under the free GR peak at 2.5 S.
C) Complex formation analysis of labeled GR-LBD* in the presence of Sgt1, Hsp90 and Aha1 by aUC sedimentation is shown. Normalized c(S) distributions were plotted against the sedimentation coefficient S [GR-LBD*: 500 nM, Hsp90: 10 μM, Sgt1: 20 μM, Aha1: 10 μM, ATP$\gamma$S: 2 mM]. Right panel: Quantification of free GR-LBD was performed by integrating the area under the free GR peak at 2.5 S.
D) Complex formation analysis of labeled Aha1* in the presence of Hsp90, Sgt1 and the Sgt1 domains TPR-CS by aUC sedimentation is shown. Normalized c(S) distributions were plotted against the sedimentation coefficient S [Aha1*: 500 nM, Hsp90: 10 μM, $Sgt1^{TPR\text{-}CS}$: 10 μM, Sgt1: 10 μM, Aha1: 10μM, ATP: 5 mM]. Right panel: Quantification of free Aha1 was performed by integrating the area under the free Aha1 peak at 2.5 S.
E) Superimposition of our crystal structure of the Hsp90-MD / Sgt1-CS complex on the structure of the Hsp90/Aha1 complex in its closed state (pdb: 6XLF) and open state (pdb: 6XLB) that shows that the binding sites of Aha1 NTD and Sgt1-CS overlap, and are incompatible with simultaneous binding.
